# Enhancers Silence Coupled Coding RNAs in Animals

**DOI:** 10.64898/2026.11.03.763931

**Authors:** Lingbo Li, Song Mei, Haolin Liu, Xinjian Liu, Qianqian Zhang, Ernesto Guccione, Haitao Li, Chuanyuan Li, Gongyi Zhang

## Abstract

The exact working mechanism and biogenesis of regulatory non-coding RNAs (rncRNAs), including enhancer RNAs (eRNAs), microRNAs (miRNAs), piRNAs, small interfering RNAs (siRNAs), transposons, and other non-coding RNAs, remain enigmatic. Conversely, the unique transcriptional regulation mechanism that differentiates animals from simple unicellular organisms remains elusive in life sciences. We find that regulatory non-coding genes are independent transcription units marked with arginine methylation of nucleosomes within their gene bodies, which is regulated by JMJD5. Enhancer RNAs are similar to those of other regulatory RNAs in targeting the 3’UTRs of coupled coding RNAs. All regulatory RNA genes share transcription factors with their target coding genes, both of which are regulated by promoter-proximal RNA Polymerase II pausing and may control the development of animals. Temporal and spatial coding genes and regulatory non-coding genes are intrinsically coupled in animals.

## Introduction

The discovery of regulatory non-coding RNAs (rncRNAs) and their role in the regulation of animal development is one of the most important events in life sciences in recent decades (Lee, Feinbaum et al. 1993, Wightman, Ha et al. 1993, Fire, Xu et al. 1998, Hamilton and Baulcombe 1999, Zamore, Tuschl et al. 2000). Compared to the well-illustrated working mechanism and biogenesis of small interfering RNAs (siRNAs)(Hamilton and Baulcombe 1999, Zamore, Tuschl et al. 2000), which was appreciated right after its discovery in life sciences, the exact working mechanism and biogenesis of microRNAs (miRNAs), piRNAs, enhancer RNAs (eRNAs), and transposons remain ambiguous, especially eRNAs. It is believed that the majority of miRNAs originate from introns (Cai, Hagedorn et al. 2004, Rodriguez, Griffiths-Jones et al. 2004), suggesting that miRNAs are the main by-products of Pol II transcription (Lee, Kim et al. 2004) and are processed by a mechanism similar to that of siRNAs (Lee, Ahn et al. 2003, Han, Lee et al. 2004). Several working mechanisms have been proposed for miRNAs, though all of them could exist or not be mutually exclusive, including blocking the synthesis of targeted protein mRNA via binding to the 3’UTR (Williams 2008), degradation of 3’ polyadenylation (Eulalio, Huntzinger et al. 2009), histone modifications (Hawkins and Morris 2008, Tan, Zhang et al. 2009), and potential cleavage of mRNA via Argonaute, similar to that of siRNA (Kawasaki and Taira 2004, Meister, Landthaler et al. 2004, Wang, Reyes et al. 2004). piRNAs were first characterized as mainly targeting transposons (Carmell, Girard et al. 2007, Gunawardane, Saito et al. 2007, Houwing, Kamminga et al. 2007, Yin and Lin 2007) via piwi proteins as platforms (Cox, Chao et al. 1998, Deng and Lin 2002), homologs of Argonaute. Several later groundbreaking discoveries showed that piRNAs originate from intergenic regions and target a large number of mRNAs during the spermgenisis of both Drosophila and mouse (Gou, Dai et al. 2014, Barckmann, Pierson et al. 2015, Goh, Falciatori et al. 2015, Watanabe, Cheng et al. 2015) as well as adult stem cells (Zhang, Kang et al. 2015), which expands our knowledge that piRNAs only originate and target transposons in the early reports. A recent excellent report from Zamore’s group showed that piRNAs may work similarly to siRNAs to degrade target mRNAs (Cecchini, Zamani et al. 2026), suggesting that all regulatory non-coding RNAs, including siRNAs, piRNAs, and miRNAs, may work similarly to degrade target mRNAs. Transposons were characterized to produce antisense RNAs (siRNAs) against themselves to prevent random translocation within the host genomes after activation (Chung, Okamura et al. 2008). However, the neglected main antiviral property of transposons as innate immunity defense tools for the hosts is starting to be appreciated recently (Hale 2022). From the definition, enhancers were characterized as enhancing the expression of their coupled coding genes (Grosschedl and Birnstiel 1980, Banerji, Rusconi et al. 1981, Benoist and Chambon 1981, Gruss, Dhar et al. 1981, Banerji, Olson et al. 1983, Gillies, Morrison et al. 1983, Mercola, Wang et al. 1983). However, this notion is challenged by a series of novel discoveries in recent decades. It was found that enhancers are independent transcription units that produce non-coding enhancer RNAs (eRNAs) (De Santa, Barozzi et al. 2010, Kim, Hemberg et al. 2010). A cluster of enhancers or enhanceosomes could form a super-enhancer (Hnisz, Abraham et al. 2013, Parker, Stitzel et al. 2013, Whyte, Orlando et al. 2013), while CDK9, BRD4, p300/CBP, Mediator, etc., were found to participate in the activities of the enhanceosome, super-enhancer, and phase-separation condensates (Tang, Vijayakumar et al. 2022). A study showed that a group of miRNAs originates from eRNAs (Xiao, Li et al. 2017, Ren, Liu et al. 2025). However, the exact function of eRNAs remains elusive, though people tend to believe that they may play some positive role in the expression of their targeted genes. Interestingly, even siRNA was found to target host mRNA in animals (Liu, Haasnoot et al. 2008). All these accumulating data suggest that some critical information on the function and generation of miRNAs, eRNAs, piRNAs, siRNAs, and other non-coding RNAs is still missing, especially intrinsic relationships with their targeted coding genes as well as among themselves.

JMJD5 is not a well-characterized orphan family member of the Jumonji protein family or the big hydroxylase family. However, knockout of JMJD5 leads to embryonic lethality at E8.5 (Oh and Janknecht 2012), which is the most severe phenotype observed from knockout among all other members, including the lysine demethylase family (Lee, Song et al. 2000), the DNA demethylase family (Moore, Le et al. 2013), the RNA demethylase family (Zheng, Dahl et al. 2013), or the hydroxylase Factor Inhibiting HIF (FIH) (Zhang, Fu et al. 2010). Only LSD1, a H3K4/9me2/1 demethylase, which is essential for the development of animals, has a similar knockout phenotype (Foster, Dovey et al. 2010). We incidentally discovered that JMJD5 has endopeptidase activity toward methylated arginine on histone subunits and continues to act as an exopeptidase after the first cut (Liu, Wang et al. 2017, Liu, Wang et al. 2018). Further characterization showed that there is a CTD of Pol II interacting domain (CID) within the N-terminus of JMJD5 (Liu, Ramachandran et al. 2020), which recognizes a unique phosphorylation pattern generated by CDK9 (Liu, Ramachandran et al. 2020). Moreover, we found that CDK9 is recruited onto the super-elongation complex (SEC) by BRD4 with the help of JMJD6, which also acts as a protease to cleave the arginine-methylated MePCE protein of the 7SK snRNP complex to release CDK9+CyclinT (Hong, Zang et al. 2010, Lee, Liu et al. 2020). Most surprisingly, we found that both BRD4 and p300/CBP work upstream of the CDK9 pathway (Liu, Li et al. 2025), and CDK9 exists only in animals, possibly functions to replace CDK10/11, and subsequently, CDK12/13, which are responsible for the generation of phosphorylated Ser2-CTD of Pol II, the elongation-active Pol II for regular genes in all eukaryotes (Liu, Li et al. 2025). ATAC-seq data showed that depletion of JMJD5 in mice led to the dramatic reduction of accessibility of transcription start sites for more than 1,371 genes (Liu, Ramachandran et al. 2020), which proved our early hypothesis that arginine-methylated nucleosomes may only exist at +1 for these Pol II pausing-regulated genes (Liu, Wang et al. 2017, Liu, Wang et al. 2018). To further explore the importance of the coupling of arginine methylation on +1 nucleosomes of the gene bodies of coding genes and Pol II pausing regulation, we carried out a more detailed analysis of arginine methylation genome-wide in human cells. To our surprise, we found that arginine methylation is ubiquitously distributed within the gene bodies of 22,022 genes (total of 30,463 peaks), including 12,284 coding genes and 9,738 non-coding genes. A separate TSS-window analysis identified 6,709 protein-coding genes in the predefined +1/+2 windows. Depletion of JMJD5 leads to dramatic accumulation of H3R2me2s on 1,240 genes, of which 349 are coding genes (**Table 1**). These data suggest that the expression of regulatory non-coding genes is mainly regulated by arginine methylation and JMJD5, while Pol II pausing regulation also participates in the regulation of non-coding genes. Detailed analysis reveals that a group of coding RNAs and non-coding RNAs are intrinsically coupled by the same group of transcription factors, which govern the development of animals and environmental cues.

### Depletion of JMJD5 leads to the accumulation of H3Rme2s within gene bodies at transcription start sites of coding genes (mRNAs)

It has been widely reported that transcription activation (Chen, Huang et al. 2000, Ma, Baumann et al. 2001, Strahl, Briggs et al. 2001, Daujat, Bauer et al. 2002, An, Kim et al. 2004) and repression (Pollack, Kotenko et al. 1999, Pal, Vishwanath et al. 2004, Guccione, Bassi et al. 2007, Scaglione, Patzig et al. 2018) (Waldmann, Izzo et al. 2011) are coupled with histone arginine methylation on histones. Arginine methylation on histone subunits is ubiquitous and essential for life in animals (Bachand 2007, Larsen, Sylvestersen et al. 2016, Rakow, Pullamsetti et al. 2020). We have proposed that these modifications may participate in the recruitment of a variety of histone modification complexes to convert inactive promoters, such as facultative repressive heterochromatin, into active docking sites via p300/CBP and MLL3/4 (de novo generation of active promoters) or vice versa to recruit repressive modification complexes to generate facultative heterochromatin (Liu, Wei et al. 2022). The arginine methylation modification should be located on the +1/N (N, an integer) nucleosomes since docking sites for long-term purposes should be at -1/N sites, which would not be affected during the multiple runs of the transcription process (Zhang 2024). One unanswered question is why there is a need to cleave these arginine-methylated histone tails, a breakdown of the hallmark of reversibility of epigenetics. One possible explanation is that these arginine-methylated histone tails are no longer recognized by histone modification complexes, which generate fragile nucleosomes along gene bodies, including pan-acetylation and pan-phosphorylation, since we found that this process is essential for Pol II to transcribe in all eukaryotes (Zhang 2024, Li, Hunter et al. 2025). Although we have solid ATAC-seq data to prove that the cleavage of arginine-methylated histone tails on +1 nucleosomes happens in MEF cells (Liu, Ramachandran et al. 2020), the concept of “tailless fragile nucleosomes” generated by JMJD5 is still not well-received in the field even after a series of stories published from our group (Hong, Zang et al. 2010, Liu, Wang et al. 2017, Liu, Wang et al. 2018, Lee, Liu et al. 2020, Liu, Lee et al. 2020, Liu, Ramachandran et al. 2020, Liu, Wei et al. 2022, Liu, Li et al. 2025). To further verify or cross-confirm our conclusion, we carried out CUT&TAG-seq on one popular arginine methylation mark on histone tails, the symmetric double arginine methylation on H3R2 (H3R2me2s), which is generated by PRMT5 and PRMT7 during transcription activation (Migliori, Muller et al. 2012) but does not happen in yeast (**Fig. 1A**), and is removed by JMJD5 (**Fig. 1B**) (Liu, Wang et al. 2017). To our surprise, there are ∼22,022 (30,463 peaks) genes (both coding and non-coding genes) having this modification in a human cancer cell line (A357) (**Fig. 1C and 1D**). Depletion of JMJD5 leads to a dramatic drop (9,131) in the overall arginine-methylated genes (**Fig. 1E**), suggesting that JMJD5 plays a critical role in the generation of arginine methylation of nucleosomes within gene bodies. Most importantly, JMJD5 knockout leads to the accumulation of these modifications of nucleosomes within the bodies of ∼1,240 genes (almost every peak represents one gene, including coding and non-coding genes) (**Fig. 1F**). These data are consistent with the results derived from ATAC-seq in MEF cells (**sFig.1**); knockout of JMJD5 led to the loss of accessibility of ∼1,481 genes at their transcription start sites, as we reported earlier (Liu, Ramachandran et al. 2020). Most importantly, this result further confirms the authentic protease activities of JMJD5, as we reported (Liu, Wang et al. 2017, Liu, Wang et al. 2018). It is beyond our expectation that there are a total of 22,022 genes with arginine methylation on nucleosomes within gene bodies. This suggests that arginine methylation on nucleosomes represents another major epigenetic modification besides lysine methylation, lysine acetylation, serine and threonine phosphorylation, and ubiquitination, which has been underappreciated in the field in the past. Detailed analysis showed that they are located within gene bodies or downstream of transcription start sites. Interestingly, we found that 6,708 coding genes (**Fig. 1G**), more than a quarter of the entire coding genes of human beings, contain arginine methylation of +1/+2 nucleosomes within their gene bodies (**Fig. 1G**), while 349 of them are affected (with an increase in the content of arginine methylation) after the knockout of JMJD5 in the cancer cell line A375 (**Fig. 1H**). These results further support our original conclusions: there exist arginine-methylated nucleosomes within gene bodies of a group of Pol II-pausing-regulated genes, and they are regulated by the protease activity of JMJD5, which generates fragile tailless nucleosomes for Pol II to transcribe through during early elongation to release paused Pol II, as pan-acetylated or pan-phosphorylated nucleosomes (Li, Hunter et al. 2025).

**Figure 1.**
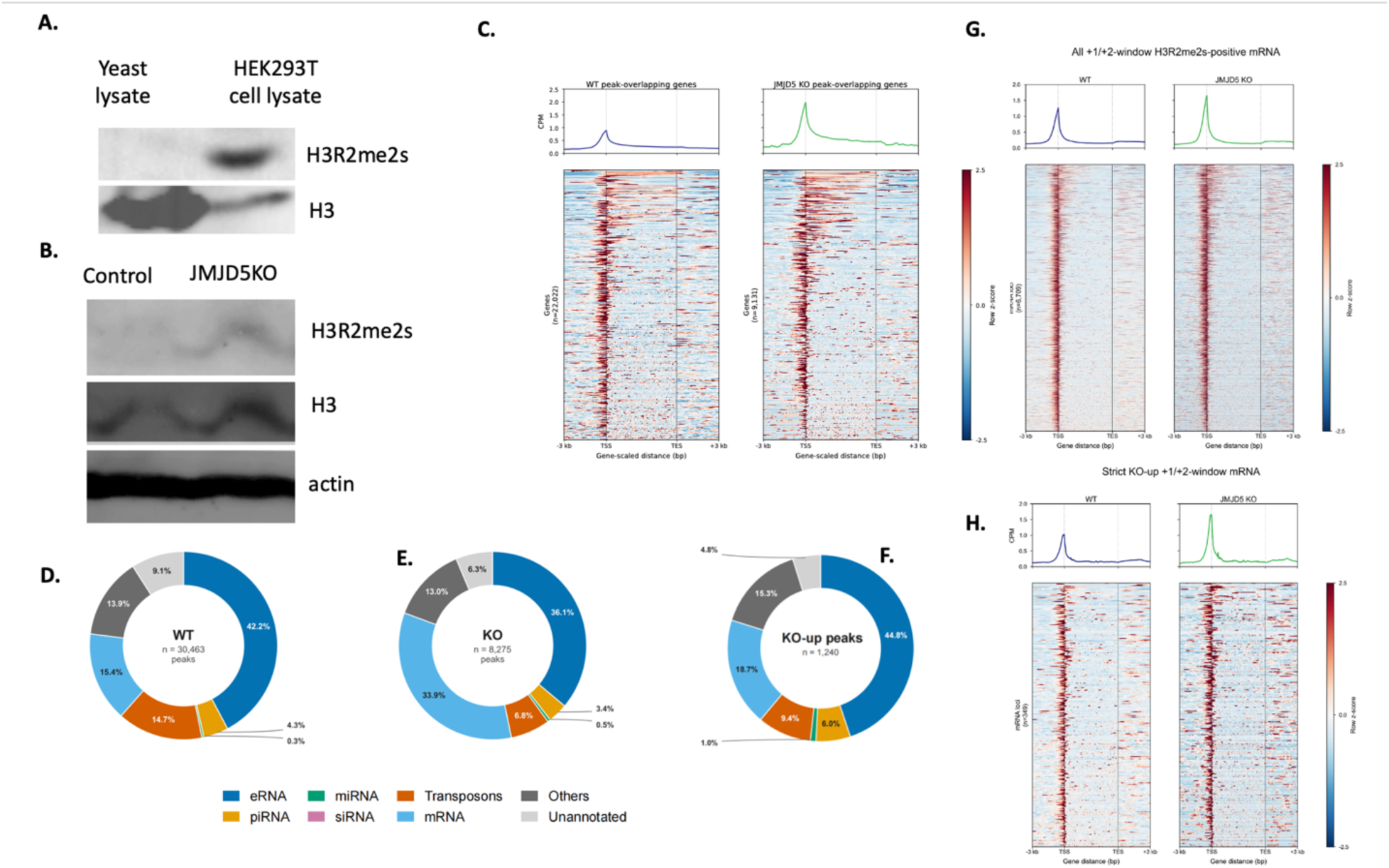
The unique histone H3R2me2s arginine methylation and ubiquitous distribution within gene bodies in animals. **A**. H3R2me2s only exists in animals but not yeast. **B**. The content of H3R2me2s increases dramatically with JMJD5 knockout in the human A375 cancer cell line. **C**. There are 22,022 genes containing these modifications in native A375, while it drops to 9,131 in JMJD5 knockout cells. **D**. A total of 30, 463 peaks in native A375. **E**. The number drops to 8,275 in JMJD5 KO. **F**. The modification increases in 1,240 genes with the knockout of JMJD5. **G**. There are 6,709 coding genes containing this modification in the +1/+2 nucleosomes. **H**. The modification increases in a total of 349 coding genes.

### Arginine methylation is ubiquitous within gene bodies of regulatory RNAs, including miRNAs, piRNAs, transposons, and other non-coding RNAs

As mentioned, there are a total of 22,022 genes containing arginine methylation within their gene bodies; besides 12,284 coding genes (dropped to 6,709 with H3R2me2s signal in +1/+2 nucleosomes after JMJD5 knockout), there are 9,738 non-coding genes containing this modification within their gene bodies. Interestingly, 46 of 122 miRNA genes are affected by the knockout of JMJD5 (**Fig. 2A, sFig. 2A**), while 390 of 5,043 H3R2me2s-marked piRNA alignment intervals (corresponding to 140 of 1,105 distinct piRNAs), are affected (**Fig. 2B, sFig. 2B**). To our surprise, 592 of 14,185 transposons are also affected by the knockout of JMJD5 (**Fig. 2C, sFig. 2C**), while 449 of 9,633 other uncharacterized non-coding genes are affected (**Fig. 2D, sFig. 2D**) (**Table 2**). As well characterized before, all these regulatory RNAs target host coding mRNAs (Lee, Feinbaum et al. 1993, Wightman, Ha et al. 1993, Fire, Xu et al. 1998, Carmell, Girard et al. 2007, Gunawardane, Saito et al. 2007, Houwing, Kamminga et al. 2007, Yin and Lin 2007). Analysis of these affected regulatory non-coding RNAs (rncRNAs), 99.5% of piRNAs are successfully matched to 3’UTRs of mRNAs (**Fig. 2E**), while 73.9% of miRNAs are matched (**Fig. 2F**). Interestingly,94.3% of transposons (**Fig. 2G**), 75.3% of others (**Fig. 2H**), and 43.3% of unannotated (**Fig.2I**) are matched too. These results suggest that most, if not all, regulatory non-coding RNA genes contain arginine-methylated nucleosomes and are regulated by JMJD5, and, most importantly, target coding mRNAs.

**Figure 2.**
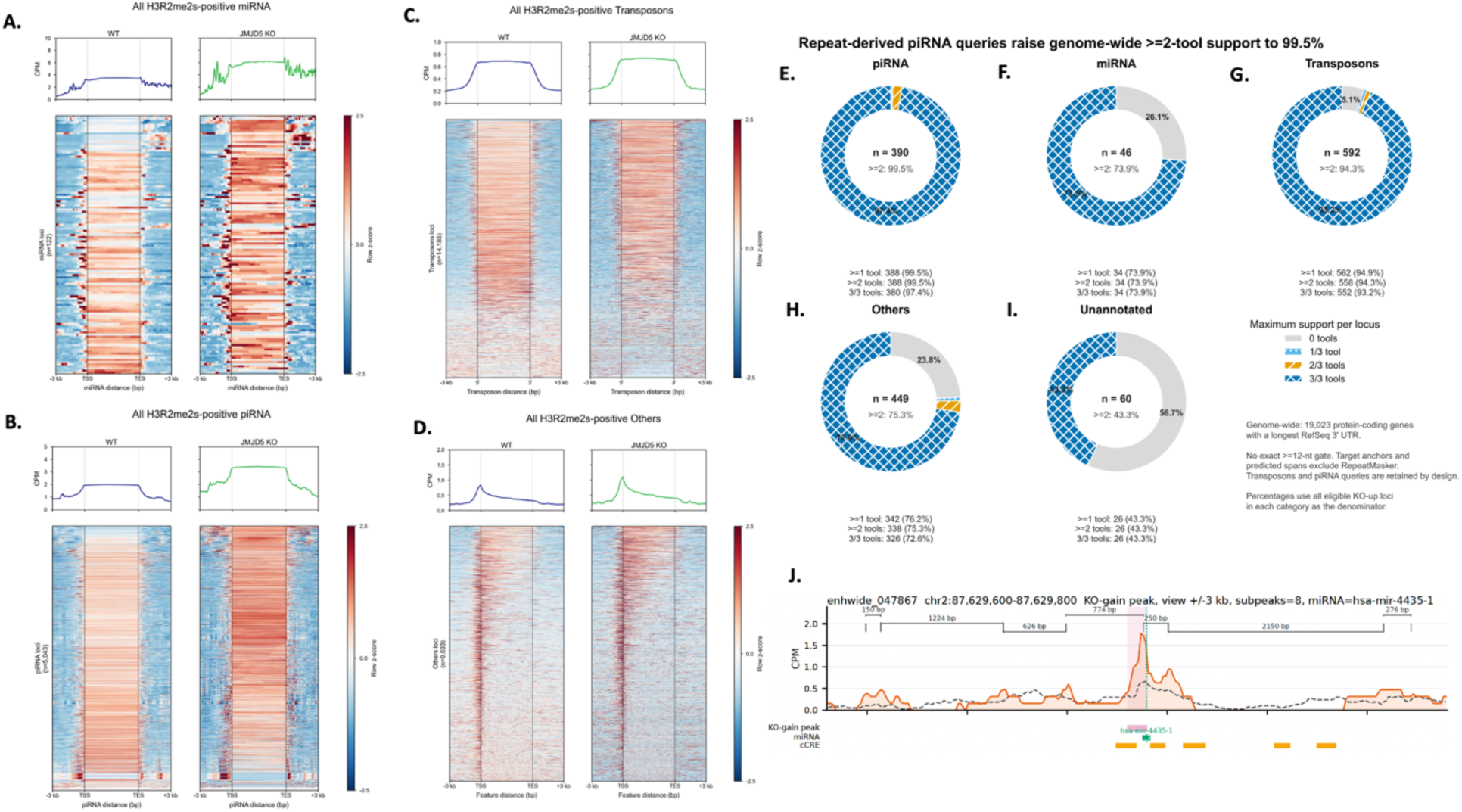
The ubiquitous existence of H3R2me2s in the bodies of regulatory non-coding genes. **A**. There are 122 miRNA genes containing this modification within their gene bodies. **B**. There are 5,043 piRNA genes containing this modification within their gene bodies. **C**. There are 14,185 transposons containing this modification within their gene bodies. **D**. There are 9,633 other non-coding genes containing this modification within their gene bodies. **E**. The modification increases in 390 piRNA bodies. **F**. The modification increases in 46 miRNA genes. **G**. The modification increases in 592 Transposons. **H**. The modification increases in 449 other non-coding genes. **I**. The modification increases in 60 unannotated genes. **J.** The modification in two nucleosomes within the gene body of has-mir-4435-1.

An interesting observation is that almost all gene bodies of rncRNA genes are occupied by arginine-methylated nucleosomes (**Fig. 2A-2D**). The length of pre-rncRNAs is variable from ∼150 to 350; it could be longer if there is a cluster of rncRNAs. A question arising here is whether these longer miRNA transcription units contain multiple arginine-methylated nucleosomes. It turns out to be true. When we inspect longer transcription units, we do see two nucleosomes showing up within the gene bodies of several miRNAs and are affected by the knockout of JMJD5 (**Fig. 2J, sFig. 2E**). This suggests that all these arginine-methylated nucleosomes are destroyed (via cleavage of histone tails by JMJD5) during transcription elongation to generate fragile nucleosomes (tailless nucleosomes) for Pol II to transcribe through for these rncRNA genes. No further complicated generation of fragile nucleosomes via pan-acetylation, pan-phosphorylation, or both is required for these rncRNA transcription units, while the process is essential for regular coding genes, as we reported (Li, Hunter et al. 2025). However, this should not be true for the coding genes (mRNAs); the bodies of these mRNA genes contain more than two nucleosomes, though we found two peaks of arginine-methylated nucleosomes in these genes as well (not shown). It should also be true that JMJD5 should cleave the histone tails of both +1 and +2 arginine-methylated nucleosomes. Our original model of the cleavage of arginine-methylated histone tails of only +1 nucleosomes should be modified or corrected (Liu, Wang et al. 2017).

### Enhancers are independent transcription units that contain arginine-methylated nucleosomes within their gene bodies

Enhancers were discovered more than four decades ago (Grosschedl and Birnstiel 1980, Banerji, Rusconi et al. 1981, Benoist and Chambon 1981, Gruss, Dhar et al. 1981, Banerji, Olson et al.

1983, Gillies, Morrison et al. 1983, Mercola, Wang et al. 1983). Not long ago, enhancers were found to be independent transcription units that produce enhancer RNAs (eRNAs) (De Santa, Barozzi et al. 2010, Kim, Hemberg et al. 2010). It was also found that eRNAs contain miRNAs (Xiao, Li et al. 2017) and were interpreted as nuclear-activating miRNAs to activate nearby genes (Liang, Zou et al. 2017), which is not consistent with the general roles of miRNAs. In this regard, the exact working mechanism and biogenesis of eRNAs remain ambiguous. To our surprise, we found that there are 14,832 enhancer genes containing arginine methylation of nucleosomes within their gene bodies (**Fig. 3A**), while 713 of 14,832 are affected after the knockout of JMJD5, with an increase in the content of arginine methylation (**Fig. 3B**). More surprisingly, most of them are matched with the 3’-UTRs of coding RNAs (**Fig. 3D**) and are located proximate to their matched mRNAs (**Fig. 3C**) (**Table 3**). These data suggest that most eRNAs, if not all, are actually antisense RNAs and could function similarly to those of other rncRNAs, including miRNAs, piRNAs, and transposons.

**Figure 3.**
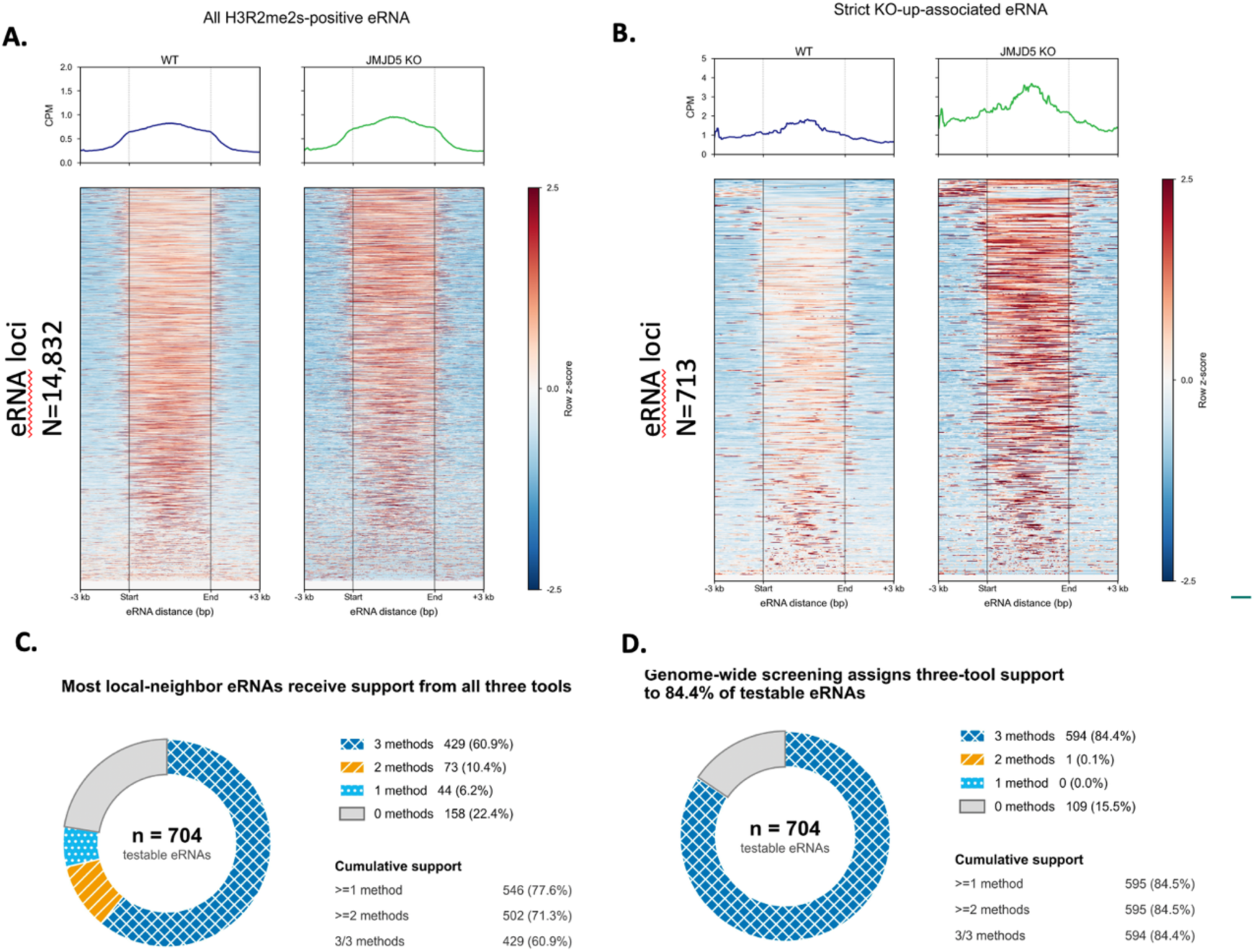
Enhancer genes also contain this modification within their gene bodies and match with 3’UTRs of mRNAs. A. There are 14,832 enhancer genes containing this modification. **B**. This modification increases in 713 eRNAs with the knockout of JMJD5. **C**. 77.6% of 704 eRNAs are located close to their target mRNAs. **D**. 84.5% of 704 eRNAs match with 3’UTRs of mRNAs genome-wide.

### eRNAs act as regulatory non-coding RNAs to target mRNA genes

It is believed that the relative distance between miRNA genes and their target messenger RNA (mRNA) genes does not rely on a fixed genomic or linear distance between their DNA loci, meaning the transcribed miRNA molecule travels through the cell, either in the nucleus or the cytosol, to bind to target mRNAs anywhere in the cell. They can be located on completely different chromosomes or far apart on the same chromosome, such as Lin-4 and LIN-14 (Lee, Feinbaum et al. 1993). However, our data show that a majority of pairs between eRNA genes and target mRNA genes are located close to each other, as we showed above (**Table 3**). A close examination of the enhancer gene, db_eRNA_12441582, and its target coding gene, the TARS3 gene, shows that db_eRNA_12441582 is located upstream of TARS3 (**Fig. 4A**, **4B**), and a characteristic match, as that of miRNAs and target mRNA genes (**Fig. 4C**, **4D**). 77.6% of 704 eRNAs are located close to their target mRNAs (**Fig. 3C**), while 84.4% of them are matched to 3’UTRs of mRNAs (**Fig. 3D**) (**Table 3**). This result suggests a preferred proximate regulation mechanism between eRNAs and mRNAs. To find out if it is also true for other rncRNAs, we examined miRNAs, piRNAs, transposons, and others. Interestingly, only 24% of piRNAs (**sFig.4A**), 54% of miRNAs (**sFig. 4B**), 64% of transposons (**sFig. 4C**), 58% of others (**sFig. 4D**), and 42% of unannotated (**sFig. 4E**) are close to their target mRNAs. A close examination of a miRNA, has-mir-941-1, which targets both the UCKL1 gene and the TPD52L2 gene, is located downstream of both coding genes UCKL1 and TPD52L2, though with opposite transcription directions (**sFig. 4F, 4G**). We found antisense sequences within has-mir-941-1 against either the UCKL1 gene (**sFig. 4H)** or the TPD52L2 gene (s**Fig. 4I**). Based on the above analysis, several common features between eRNAs and other rncRNAs could be derived. First, eRNA genes have similar compositions of arginine-methylated nucleosomes in their gene bodies (**Fig. 3A**, **3B**). Second, eRNA genes are mostly located adjacent to their target mRNAs (**Fig. 3C**). Third, we found that enhancers/enhanceosomes/super-enhancers are coupled with Pol II pausing regulation via BRD4 and p300/CBP (Liu 2025), while miRNA genes are rich within these super-enhancer clusters (Hnisz, Abraham et al. 2013, Parker, Stitzel et al. 2013, Whyte, Orlando et al. 2013), suggesting overlapping between eRNAs and miRNAs. We do find several miRNAs in the library of eRNAs in our data, supporting the earlier discoveries from others (Xiao, Li et al. 2017). Most importantly, eRNAs are similar to other rncRNAs that target mRNAs via the 3’UTRs. Based on the analysis, it may be safe to claim that eRNAs actually work like other regulatory RNAs to silence their target coding genes (mRNAs) in the cytosol; instead of the original definition of enhancers in the nucleus, there are actually silencers that degrade mRNAs in the cytosol when the missions of the latter are accomplished during development and acute responses.

**Figure 4.**
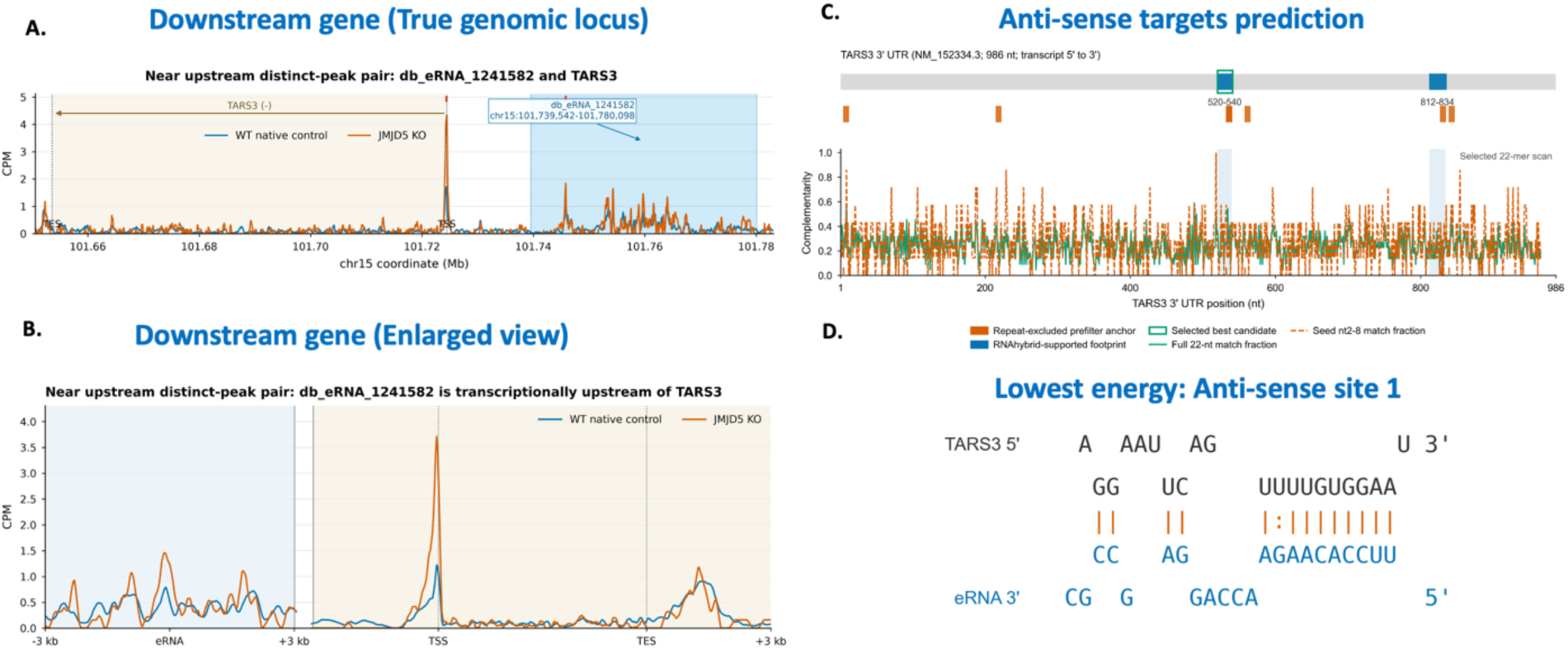
eRNAs work similarly to miRNAs to target mRNAs with a characteristic miRNA match mode; enhancers are silencers. **A.** db_eRNA_124582 and its target TARS3. **B**. Enlarged view of the pair. **C**. Sequence matching analysis. **D**. The match site.

### rncRNA genes and eRNA genes are regulated by the same group of transcription factors as their target coding genes

Currently, people believe that miRNAs mostly originate from introns and are generated by Pol II (Cai, Hagedorn et al. 2004, Lee, Kim et al. 2004, Rodriguez, Griffiths-Jones et al. 2004). An excellent report from Dr. Richard Young’s group showed that promoter regions of a group of miRNAs within a super-enhancer have multiple transcription factor binding sites (Whyte, Orlando et al. 2013), similar to those of Pol II-pausing-regulated genes (Liu 2025, Zhang 2026). One unanswered question from these observations is whether miRNAs are actually independent transcription units, if not all of them, since these transcription factors are enough to trigger the activation of coupled genes, as elegantly demonstrated by Dr. Maniatis and others’ groups (Giese, Kingsley et al. 1995, Thanos and Maniatis 1995). Furthermore, it was also found that Mediators, CDK9, BRD4, and p300/CBP, are present within the super-enhancer cluster (Tang, Vijayakumar et al. 2022), suggesting that these miRNAs are independent transcription units having a similar transcription activation mechanism as that of Pol II pausing-regulated genes (Liu 2025, Zhang 2026). We wonder whether these miRNAs may have similar transcription factors, which bind to their promoter regions to trigger the expression of both miRNAs and mRNAs. Sequence analysis of the promoter regions of both hsa-miR-941 and UCKL1 via the JASPAR program, we do find similar transcription factor binding sites (**sFig. 5A**) as well as TPD52L2 (**sFig. 5B**) (**Table S1**).

It was well established that promoters of eRNA genes also contain binding sites for all four major transcription factors, similar to those of miRNA genes (Giese, Kingsley et al. 1995, Thanos and Maniatis 1995). Most importantly, people found that there exist Mediator, BRD4, and p300/CBP at active eRNA genes within the super-enhancer (Tang, Vijayakumar et al. 2022). To our satisfaction, upon detailed examination of promoters of eRNAs and their target mRNAs, we found a similar transcription factor binding mode as those of miRNA mentioned above (**Fig. 5**). We found exactly the same group of transcription factor binding sites for the promoter of the db_eRNA_12441582 enhancer gene (**Fig. 5A**) and the promoter of TARS3 gene (**Fig. 5B**), including wTHT family members, SPIB and Spi1, the bZIP family members, TFAP2B and BHLHE22, the Ig family member, Arnt, and a large group of Zinc-finger members, including NR6A1 (**Table 4**). This discovery suggests that eRNA genes are activated by a similar group of transcription factors as coupled coding genes and are transcribed independently, as those of miRNA genes.

**Figure 5.**
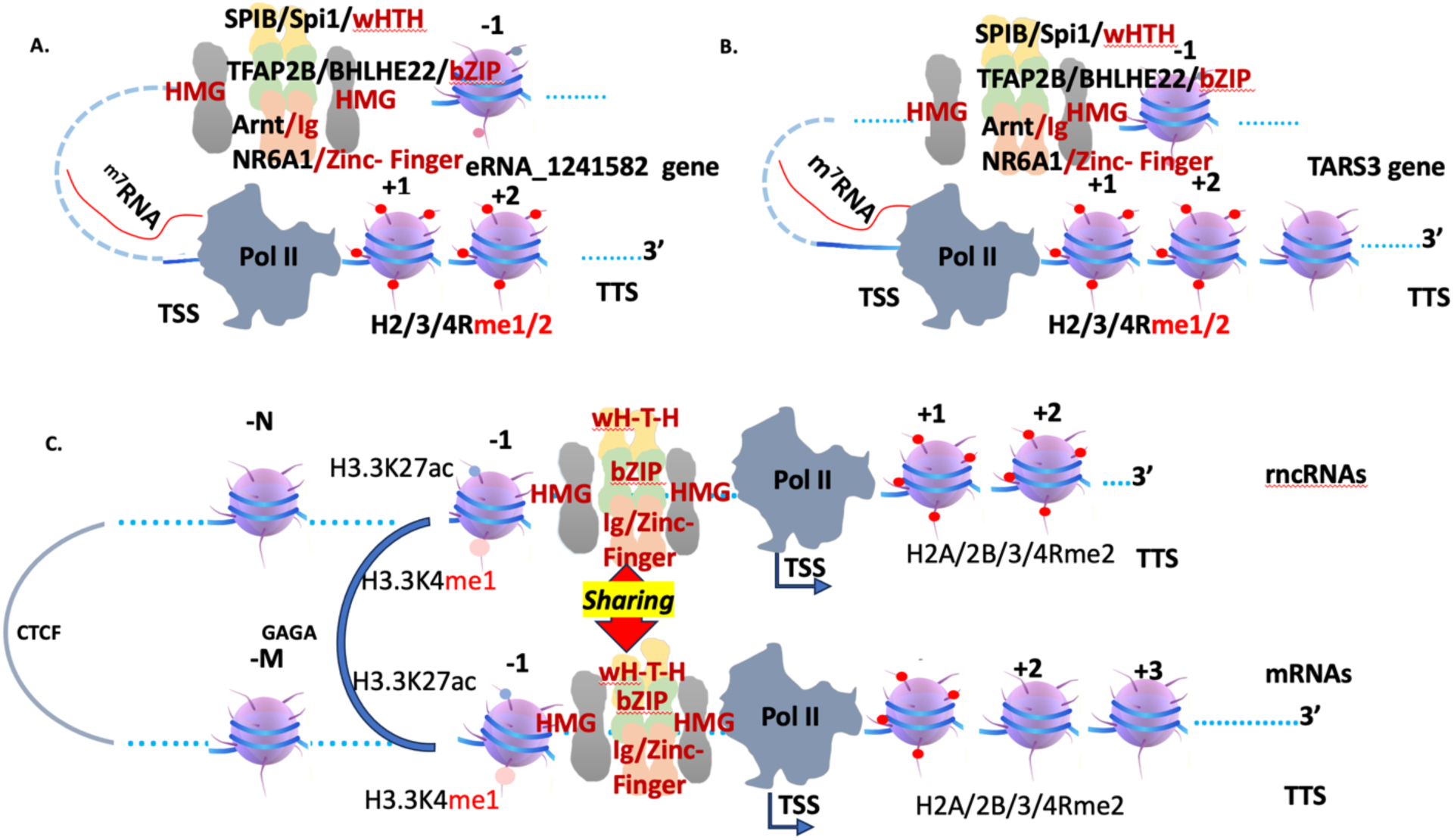
Both regulatory non-coding genes (rncRNAs) and their target genes share the same group of transcription factors during activation. **A.** Four transcription factor family members, SPIB/Spi1-wHTH, TFAP2/BHLHE22-bZIP, Arnt-Ig family, and NR6A1-Zinc-finger, potentially bind to the promoter of the eRNA_1241582 gene. **B.** A similar group of transcription factors potentially binds to the promoter of the TARS3 gene. **C.** A model of transcription factor sharing between rncRNAs and mRNAs, which could be brought together by GAGA and CTCF tethering proteins.

Based on these data, we claim that most, if not all, rncRNA genes and eRNA genes (**Fig. 5A**) share transcription activation factors with their target mRNAs (**Fig. 5B**). It is well established that looping and tethering proteins such as GAGA (Li, Tang et al. 2023) and CTCF (Batut, Bing et al. 2022) could bring long-distance DNA together. We propose that promoters of rncRNAs/eRNAs and mRNAs could be brought together to form a cluster of transcription hubs (super-enhancer clusters); a model could be derived (**Fig. 5C**). In this model, both rncRNAs/eRNAs and mRNAs share the same group of transcription factors, including a high mobility group (HMG) family member, a Winged-Helix-Turn-Helix (wHTH) member, a basic leucine zipper (bZIP) family member, and an Ig-domain or Zinc-finger family member, similar to that of the enhanceosome reported earlier (Giese, Kingsley et al. 1995, Thanos and Maniatis 1995). It was reported that multiple miRNA genes could form a cluster and super-enhancers (Hnisz, Abraham et al. 2013). If we bring these rncRNA genes and their coupled mRNA genes into consideration, a cluster model of multiple rncRNA genes and multiple mRNA genes could be derived (**Fig. 6**). This model may interpret the 3D topologically associated domains (TADs) of the entire animal genomes (Dixon, Selvaraj et al. 2012, Nora, Lajoie et al. 2012).

**Figure 6.**
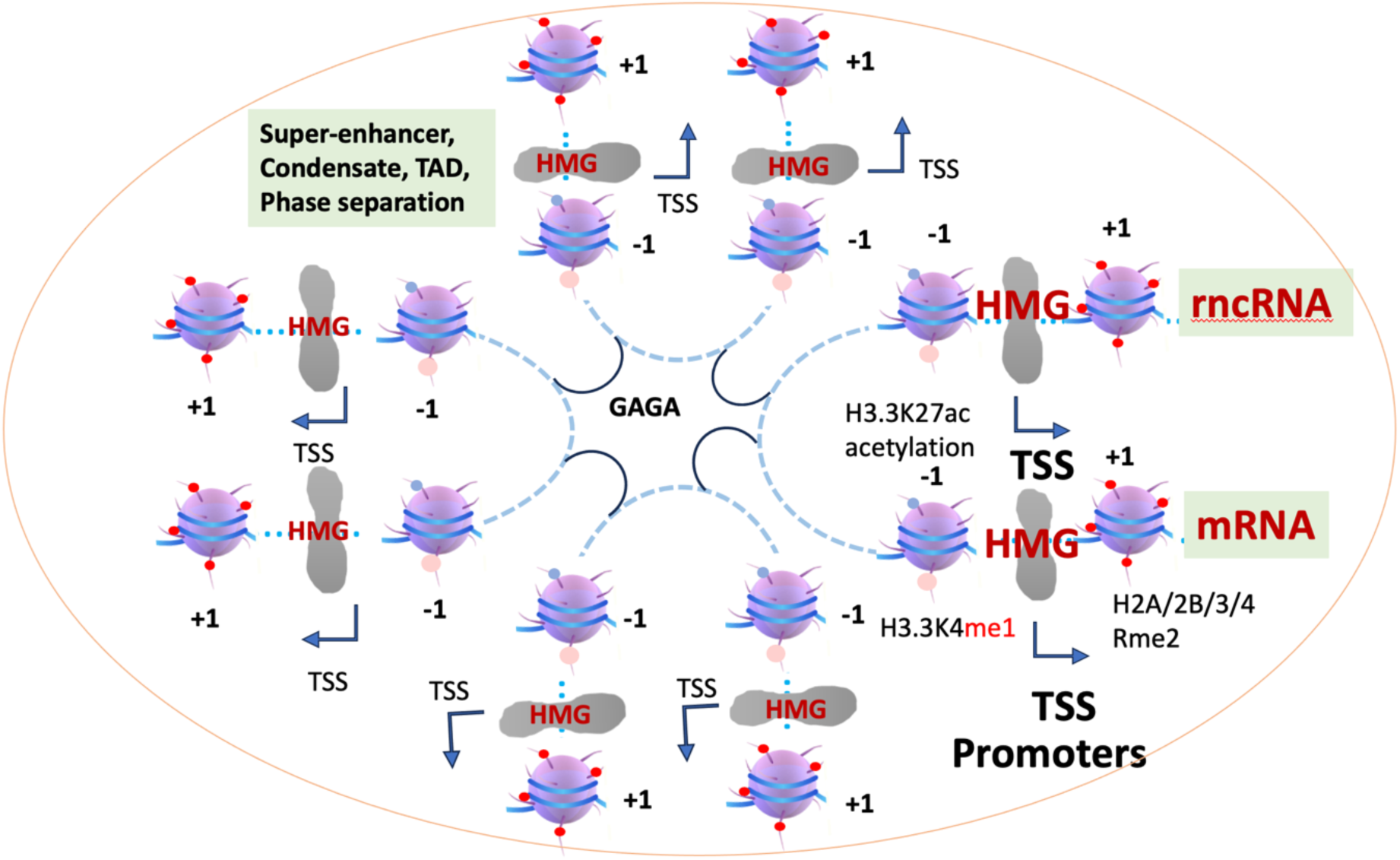
A model of a cluster of pairs of rncRNA genes and their target mRNA genes. These clusters (super-enhancers) may build up these phase-separated condensates or topologically associated domains (TADs) within the nucleus.

## Discussion

Arginine methylation is ubiquitous and essential in animals (Larsen, Sylvestersen et al. 2016). A series of studies showed that arginine methylation on histone subunits of nucleosomes is required for both transcription activation and repression (Liu, Wei et al. 2022). It remains a long-standing question of how this arginine methylation on nucleosomes participates in the regulation of gene transcription activities. We proposed that arginine methylation on the +1 nucleosome of a group of Pol II-pausing regulated genes may generate docking sites for co-activators or co-repressors, respectively (Liu, Wei et al. 2022). It is a complete surprise for us to find out that arginine methylation happens on the nucleosomes along gene bodies of regulatory non-coding RNA genes and is removed by JMJD5 to generate fragile tailless nucleosomes for Pol II to transcribe and generate pre-rncRNAs during transcription activation. This groundbreaking discovery shows that histone arginine methylation is not only ubiquitous but also essential to build up unique gene bodies of nucleosomes, which is required for two major functions: a docking site to recruit activators or repressors and histone tail cleavage targets for JMJD5/JMJD7 to generate fragile tailless nucleosomes in rncRNA genes. Furthermore, this modification stands out as one of the major epigenetic modifications, besides lysine methylation, lysine acetylation, Serine/Theorine phosphorylation, and ubiquitination in animals. More surprisingly, instead of the hallmark of reversibility of these established modifications via corresponding demethylases, deacetylases, deubiquitinases, and phosphatases, arginine methylation is removed by complete destruction of histone subunits by the protease activities of JMJD5 and possibly JMJD7(Liu, Wang et al. 2017, Liu, Wang et al. 2018). It also suggests that tailless nucleosomes are a ubiquitous form of fragile nucleosomes in animals, similar to those of pan-acetylation and pan-phosphorylation, as we discovered in eukaryotes (Li, Hunter et al. 2025). It is safe to claim that the exact function and working mechanism of histone arginine methylation is finally resolved.

It is an underappreciated topic regarding the transcriptional regulation of all rncRNA genes. It is still ambiguous whether there exist independent transcription units for these rncRNA genes since people believe that the majority of miRNAs are derived from introns of coding genes or byproducts of mRNAs; there is no need for independent transcription regulation, though it was found that enhancers seem to act as independent transcription units (De Santa, Barozzi et al. 2010, Kim, Hemberg et al. 2010), as well as transposons (Chung, Okamura et al. 2008). Here, we show that most rncRNA genes, if not all, are independent transcription units and are activated by a group of transcription factors, similar to that of coupled coding genes. Furthermore, fragile nucleosomes (tailless here) along gene bodies are essential for Pol II to transcribe along gene bodies, similar to the role of fragile nucleosomes of hyperacetylation or phosphorylation in regular coding genes. However, due to the instant processing of these transcribed non-coding genes, they do not require polyadenylation at their 3’-end and are terminated by the Integrator complex, which helps us to speculate that rncRNA-coupled mRNAs may be terminated in a similar form by Integrator without 3’ polyadenylation, since they share transcription initiation and elongation as well as a short lifespan for these Pol II pausing-regulated genes (Zhang 2026). A model of the unique transcription mechanism in animals, including the CDK9-dependent generation of active Pol II (bypassing CDK10/11/ and CDK12/13), the NELF-dependent elongation (instead of Paf1c), and the Integrator-dependent termination (instead of cleavage and polyadenylation stimulating factor, CPSF), has been proposed (Zhang 2026).

The discovery that enhancers express small non-coding RNAs (De Santa, Barozzi et al. 2010, Kim, Hemberg et al. 2010) raised a major puzzle regarding the exact function and working mechanisms of these eRNAs. Later on, people found that eRNAs also contain miRNAs (Xiao, Li et al. 2017) and assigned a possible function in vivo (Liang, Zou et al. 2017). However, the exact function of enhancers remains elusive, though they are supposed to enhance the activities of coupled genes from the original definition. It is a big surprise that we find that most of the 704 pieces of affected eRNAs after JMJD5 knockout match 3’UTRs of mRNAs. This result suggests that eRNAs act similarly to other rncRNAs to target mRNAs when needed. The function of eRNAs is finally revealed. Most importantly, enhancers are actually silencers that degrade coupled mRNAs in the cytosol, instead of regulating the transcription activities of target coding genes in the nucleus.

The discoveries of the enhanceosome did not trigger further attention after the first wave of bursts (Giese, Kingsley et al. 1995, Thanos and Maniatis 1995). The discoveries of super-enhancers brought back this unique transcription activation mechanism into hot frontiers (Hnisz, Abraham et al. 2013, Parker, Stitzel et al. 2013). However, one major feature of this regulation is still underappreciated: the requirement of four or five major transcription groups that trigger the activation of a targeted gene, for example, type-I beta after viral infection. We propose that these four or five groups of transcription factors could be categorized into four to five major transcription factor families, each with a specific function besides recruiting p300/CBP. The high mobility group family (HMG), which is constitutively associated with genomes, could play a pioneering-factor role in generating nucleosome-free regions (NFRs) at the promoter regions of all genes during the stage of DNA replication in the cell cycle. The exact function of other family members remains ambiguous. However, it should be safe to claim that these four or five groups of transcription factors are both necessary and sufficient to trigger the activation of a transcription unit, either coding genes or non-coding genes. Bearing this feature in mind, we find that all rncRNAs and their coupled mRNAs share the same group of transcription factors and could be activated at the same time. It should not be difficult to understand that this coupling between rncRNAs and mRNAs could be crucial to determine the life spans of these temporal and spatial mRNAs during development or in response to environmental cues (Zhang 2026). However, how these rncRNAs are activated to start degrading target mRNAs when the mission of the latter has been accomplished should be an interesting topic for future investigation.

One unresolved question in the field is how these rncRNAs degrade their coupled coding mRNAs since the matching sequences between these rncRNAs and mRNAs are shorter than those of siRNA and their target mRNAs. One possible explanation is that these short-lived mRNAs contain 3’UTRs without polyadenylation, which may be enough for rncRNAs to trigger complete degradation with the help of exonucleases after first recognition of mRNAs by rncRNA-loaded Ago homologs. The similar working mechanism between siRNAs and piRNAs, which was revealed recently (Cecchini, Zamani et al. 2026), may support this speculation. Further characterization is required.

Finally, the exact functions and working mechanisms of siRNAs and transposons have been well characterized. miRNAs, piRNAs, eRNAs, and other uncharacterized or unannotated non-coding RNAs are all found to target mRNAs, with similar transcriptional expression mechanisms and similar gene unit building. There is no obvious criterion to distinguish them. Furthermore, they all use Ago/Argonaute protein family members or piwi family members to build up the universal RNA-induced silencing complexes (RISCs) to target mRNAs. Most importantly, miRNAs, piRNAs, and eRNAs are all found to play essential roles in the development of animals. As we know, a large group of genes is expressed temporally and spatially during the development of animals as well as during the responses to environmental cues; there must exist a mechanism to remove these short-lived mRNAs when their missions are accomplished (Zhang 2026). We claim that all these rncRNAs are born to digest these short-lived mRNAs. We may put them all together as one group of rncRNAs. In this regard, the intrinsic coupling between short-lived mRNAs and rncRNAs is established, which not only governs the development and acute responses in animals but also may be the key to differentiating animals from simple unicellular organisms (Zhang 2026).

In conclusion, during development or in response to environmental cues, a group of genes is temporally and spatially expressed. Most rncRNAs, if not all, are born to destroy their target mRNAs when their missions are accomplished, are expressed simultaneously with their target mRNAs, are activated by the same group of transcription factors as their target mRNA genes, and are regulated by the promoter-proximal Pol II pausing mechanism (Zhang 2026).

## Material and Methods

### The generation of JMJD5 knockout in A375 cancer cell line

JMJD5 knockout cell lines were generated using lentivirus-mediated CRISPR/Cas9 technology. Single-guide RNA (sgRNA) sequences (CACCG GGCCGTGCATGCAGAAGTGG; AAAC CCACTTCTGCATGCACGGCC C) targeting the JMJD5 gene were annealed and ligated into the lentiCRISPRv2 (deposited by Dr. Feng Zhang of MIT to Addgene, Cambridge, MA) at the BsmB1 site, which co-expresses Cas9 and sgRNA in the same vector. The sgRNA-encoding CRISPR lentivirus vector plasmids were transduced into 293T cells together with second-generation packaging plasmids (psPAX2, pMD2.G; both deposited at Addgene by Dr. Didier Trono of EPFL (École Polytechnique Fédérale de Lausanne, Switzerland) following a published, calcium phosphate-based Trono Lab protocol (http://tronolab.epfl.ch/page58122.html). Supernatants containing lentivirus particles were collected 48 hours after transfection. To delete the JMJD5 gene, A375 human melanoma cells were infected with lentivirus carrying sgRNA and selected with 1 µg/ml puromycin for 10 days. Cells were then seeded at super-low density to allow colony formation from a single cell. Colonies were picked and expanded for knockout validation by Western blot. The primary antibody for JMJD5 detection was used from Abcam (ab106391) and Santa Cruz Biotechnology(sc-377078).

### Western blots on yeast samples and A357 cell lines with and without JMJD5

Human HEK293 cell lysate was prepared in the RIPA lysis buffer, and the yeast cell lysate was prepared in the yeast whole cell lysis buffer (Zymo Research). Equal amount of lysate was loaded on the SDS gel and transferred to the PVDF membrane. H3R2me2s antibody (Millipore #07585) was probed, stripped, and H3 antibody (Abcam #ab1791) was subsequently probed for the blotting detection.

### CUT&Tag-seq data preparation

CUT&Tag-seq was performed according to the published method (Kaya-Okur, Wu et al. 2019). Briefly, ∼100,000 A375 cells, with and without JMJD5, were spun down and washed once with wash buffer. The cells were resuspended in 1.5 mL wash buffer and then bound to Concanavalin A-coated beads. The cells with beads were resuspended in 800 µL ice-cold antibody buffer containing H3R2me2s antibody, a gift from Dr. Ernesto Guccione’s group (Migliori, Muller et al. 2012), then placed on a nutator at 4 °C and incubated overnight. After washing, resuspended the cells with beads in Dig-wash buffer with secondary antibody from Antibodies-online (ABIN102153, and placed the tubes on a nutator at room temperature for 30–60 min. Then washed with 0.8 - 1 mL Dig-wash buffer three times. Next, the cells with beads were resuspended in Dig-300 buffer mix with pA-Tn5 adapter complex (C01070001, Diagenode, NJ). Place the tubes on a nutator at room temperature for 1 hr. After tagmentation, the DNA was extracted. Purified DNA was amplified using the following conditions: Cycle 1: 72 °C for 5 min; Cycle 2: 98 °C for 30 sec; Cycle 3: 98 °C for 10 sec; Cycle 4: 63 °C for 10 sec; Repeat Cycles 3-4 13 times; 72°C for 1 min and hold at 4 °C. The amplified libraries were purified and size-selected, and the quality and quantity of libraries were assessed on 4150 TapeStation System (Agilent, CA). The pair-end sequencing of DNA libraries was performed on an Illumina NovaSEQ6000 platform.

### Preliminary CUT&Tag-seq data processing

Raw sequencing reads were trimmed to remove adaptors and low-quality sequences using TrimmomaticPE (version 0.33). The trimmed sequencing reads were aligned to the hg38 reference genome using Bowtie2 (version 2.3.4.1) with very-sensitive and -x 2000 parameters. The read alignments were filtered using SAMtools (version 1.7) to remove PCR duplicates. Narrow peaks were called by MACS2 (version 2.1.2) with the q-value cut-off of 0.05. To analyze loss-of-binding sites after JMJD5 knockout, CUT&Tag-seq reads of A357 samples were compared with those of A357 samples with JMJD5 knockout using the R package DiffBind (version 3.12.0) (Ross-Innes, Stark et al. 2012). All raw data were processed by Basepair (https://app.basepairtech.com/). Bam files from Basepair were used in DeepTools of Galaxy (https://usegalaxy.eu/) to generate bamCompare files, computeMatricx files, and final heatmap figures (Ramirez, Ryan et al. 2016).

### Identification of candidate H3R2me2s intervals

The 24 GRCh38 primary chromosomes were divided into non-overlapping 200-bp bins. For library *s*, the expected bin count was λ*_s_* = *N_s_* × 200/*G*, where *N_s_* is the number of usable fragments in that library and *G* is the combined length of the primary chromosomes. An upper-tail Poisson *P* value was calculated for each library, and the value from the library with the larger raw bin count was used for screening. Chromosome *c*, containing *n_c_* bins, was allocated α/24 of the genome-wide error budget, which was Bonferroni-distributed across its bins. Bins were retained when max(*c*_WT_, *c*_KO_) ≥ 12 and *P* ≤ α/(24*n_c_*), with α = 1 × 10^−4^. If fewer than ten bins passed on a chromosome, Benjamini–Hochberg adjustment was applied across that chromosome, using adjusted *P* ≤ 1 × 10^−4^ and the same count threshold. Selected bins separated by no more than two intervening bins were merged into candidate intervals.

### Quantification of H3R2me2s differences between libraries

Fragments were counted in each candidate interval for both libraries. Counts per million usable fragments (CPM) were calculated as count/*N* × 10^6^, where *N* is the usable-fragment total for the corresponding library. The normalized fold change was calculated as

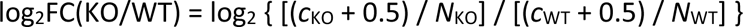

where *c* is the interval fragment count and *N* the usable-fragment total for the indicated library. For each interval, a two-sided Fisher’s exact test compared interval and non-interval fragment counts between the KO and WT libraries, and *P* values were adjusted across candidate intervals by the Benjamini–Hochberg procedure to control the false discovery rate (FDR). An interval was designated affected when the FDR was ≤ 0.05, |log_2_FC(KO/WT)| was ≥ 1, and at least 20 fragments were observed in either library.

### Genome annotation and feature classes

All analyses used the GRCh38/hg38 assembly. Protein-coding genes and representative transcription start sites (TSSs) were obtained from the GENCODE v50 primary-assembly annotation (Ensembl 116). When a single transcript per gene was required, protein-coding transcripts were prioritized in the order MANE Select, Ensembl canonical, APPRIS principal, GENCODE basic, and then all remaining transcripts. Precursor miRNA loci were obtained from UCSC GRCh38/hg38 wgRna (miRBase Release 22, March 2018) and supplemented with non-overlapping GENCODE v50 miRNA loci. piRNA loci were obtained from piRNAdb v1.7.6 hg38 and deduplicated by coordinate and strand. Transposable-element intervals were obtained from the UCSC hg38 RepeatMasker rmsk table and restricted to the LINE, SINE, LTR, DNA, Retroposon, and rolling-circle classes. Other non-coding RNA loci were GENCODE v50 genes whose gene_type was neither protein_coding nor miRNA.

A database-defined eRNA reference was assembled from FANTOM5 transcribed enhancers (Andersson et al., 2014), HACER (Wang et al., 2019), and eRNAbase (Song et al., 2024). FANTOM5 and eRNAbase provided GRCh38 coordinates, whereas HACER hg19 records were converted to GRCh38 with UCSC liftOver (-minMatch=0.95). Records with invalid coordinates or on non-primary chromosomes were removed, exact within-source duplicates were collapsed, and loci overlapping across sources were merged under a zero-gap rule while retaining provenance.

Candidate intervals were annotated by direct genomic overlap and by the nearest GENCODE gene. A feature was considered H3R2me2s-positive when it overlapped a candidate interval by at least 1 bp. The same Poisson-bin procedure was also applied independently to the WT and KO libraries. Peak-overlapping genes were defined by at least 1 bp of overlap between a peak and a GENCODE v50 gene, and were deduplicated by gene_id.

For mutually exclusive peak-level classification, each independent WT peak, independent KO peak, or strict KO-up interval was assigned to the first matching class in the priority order miRNA, piRNA, siRNA, eRNA, transposable element, mRNA, other non-coding RNA, and unannotated; peaks without a matched annotation were classified as unannotated. A protein-coding gene was retained in the +1/+2 set when the center of a significant 200-bp component bin fell within the strand-aware +1 (+40 to <+190 bp) or +2 (+190 to <+360 bp) window downstream of its representative TSS. Strict KO-up intervals were assigned to the nearest GENCODE gene: for intervals overlapping multiple genes, the gene with the greatest overlap was selected; otherwise, distance was measured from the interval midpoint to the nearest gene boundary. The +1/+2 and strict nearest-gene sets were then intersected by gene ID.

### Metagene profiles and heat maps

Signal profiles and heat maps were generated from CPM-normalized bigWig tracks using deepTools v3.5.6. computeMatrix scale-regions was run with 3-kb flanks, a 5-kb scaled feature body, and 25-bp bins, treating missing data as zero and preserving the input region order. Strand-aware features were plotted from TSS to TES, whereas unstranded eRNAs and peak intervals were plotted from genomic start to end. Mean profiles used raw CPM values. Feature-category, eRNA, and +1/+2 mRNA heat maps were z-standardized by row across WT and KO using the population standard deviation (ddof = 0); values were clipped to the range −2.5 to +2.5, and rows were sorted by descending raw mean across both libraries. Heat maps of independent WT and KO peak-overlapping genes used library-specific gene sets, row scaling, and ordering.

### RNA–mRNA 3ʹ UTR complementarity and enhancer-associated miRNA analyses

Protein-coding targets were defined by intersecting GENCODE protein-coding genes with UCSC hg38 ncbiRefSeq. The longest 3ʹ untranslated region (UTR) was selected per gene, with curated NM_ transcripts preferred for length ties, and exons were joined in transcript orientation. This reference was used for both screens below.

Non-eRNA queries were constructed from piRNA, precursor miRNA, transposable-element, other non-coding RNA, and unannotated features. Query cores were the feature–interval overlap for piRNA, transposable elements, and directly overlapping other non-coding loci; the 22-nt segment at the closest gene boundary for nearest-assigned other non-coding genes without overlap; the complete precursor locus for miRNA; and the strict KO-up interval itself for unannotated features. KO-gain miRNA loci were selected by counting fragments over each precursor locus with 2-kb flanks, comparing precursor-window with non-window counts between KO and WT by two-sided Fisher’s exact test with Benjamini–Hochberg adjustment, and retaining loci at FDR ≤ 0.05, log_2_FC(KO/WT) ≥ 1 (0.5-fragment pseudocount), and max(*c*_WT_, *c*_KO_) ≥ 10. eRNA query cores were the intersections of eRNA annotations with overlapping strict KO-up intervals, extended within the annotated eRNA boundaries when shorter than 22 nt. Because the eRNA annotations were unstranded, both transcript orientations were tested. In both screens, candidate 22-nt windows required 20–80% GC content, Shannon entropy ≥ 1.45 bits, and a maximum homopolymer length of 6 nt; features and loci without a retained window were assigned no predictor support.

The two screens differed in their prefilters. Genome-wide non-eRNA candidates required the reverse complement of query nucleotides 2–12 as an 11-nt anchor; up to two sites per feature– gene pair were retained, ranked by exact-complement extension, lower anchor prevalence, query entropy, and GC balance, and evaluated in cumulative batches of 100, 1,000, 10,000, and all remaining sites until a three-predictor-supported site was found or the candidates were exhausted. The local eRNA screen included the nearest flanking gene on each side and all overlapping genes, required a reverse-complementary 7mer-m8 match to query nucleotides 2– 8, and retained up to five windows per eRNA-orientation–gene pair; the genome-wide eRNA screen used a 16-nt anchor spanning query nucleotides 2–17, retained one site per pair, and excluded anchors present in the 3ʹ UTRs of more than 100 genes.

All prefiltered candidates were evaluated with RNAduplex v2.5.1, RNAhybrid v2.1.2, and miRanda v1.9, using identical settings in both screens. Each predictor required a minimum free energy (MFE) ≤ −20 kcal mol^−1^ and a predicted duplex overlapping the prefiltered site; RNAduplex and miRanda additionally required coverage of query nucleotides 2–8, and miRanda a score ≥ 140. RNAhybrid was run with -c -d 1,1 -b 5 -f 2,7 -e -20 -m 100000 -n 30, which constrains the seed helix to query nucleotides 2–7, and miRanda with -sc 140 -en -20 -quiet. Candidates were discarded when the target anchor or a method-specific target span overlapped UCSC hg38 RepeatMasker annotations; query-side repeat overlap was likewise excluded, except for piRNA and transposable-element queries. Support was summarized per feature or locus as ≥ 1, ≥ 2, or 3-of-3 predictors.

Each complexity-filtered eRNA query window was independently subjected to 100 mononucleotide permutations that preserved nucleotide composition, with unchanged sequences redrawn up to 20 times. Every replicate was rescanned through both eRNA prefilters. Enrichment was summarized by the observed-to-null mean ratio, *z* = (observed − null mean)/null SD, and the upper-tail add-one empirical *P* value, *P* = [1 + number(null ≥ observed)]/(*R* + 1).

Precursor miRNA loci were classified as enhancer-overlapping when they overlapped a proximal (pELS) or distal (dELS) enhancer-like candidate *cis*-regulatory element from ENCODE SCREEN v4. The candidate downstream gene was the nearest representative TSS in the direction of miRNA transcription, regardless of target-gene strand. Mature miRNA–mRNA relationships were queried in TargetScanHuman 8.0, miRDB v6.0, and miRTarBase v10.0, with additional support from ENCORI/starBase AGO cross-linking immunoprecipitation sites (clipExpNum ≥ 1), miRWalk human 3ʹ UTR predictions, and *de novo* RNAhybrid and miRanda predictions run with the settings above.

### Promoter motif analysis

For *UCKL1*, the region from −500 to −50 bp relative to the MANE Select TSS (ENST00000354216.11) was extracted from GRCh38 and scanned on both strands with MOODS v1.9.4.1 against the JASPAR 2026 CORE vertebrates non-redundant collection, using a uniform nucleotide background, a pseudocount of 1 × 10^−4^, and a per-site *P*-value threshold of 1 × 10^−4^. The same settings were applied to the eRNA–mRNA pair analysis, in which two regions were scanned: the strict KO-up interval overlapping db_eRNA_1241582 (chr15:101,745,600– 101,746,000; peak_022768) and the strand-aware *TARS3* promoter, extending from −2,000 to +40 bp relative to the MANE Select TSS (chr15:101,724,432–101,726,472). Unique transcription-factor names were ranked by their best relative PWM score and then by the score margin above the motif-specific threshold.

## Acknowledgments

We thank National Jewish Health and the University of Colorado Anschutz Medical Campus for providing the collaborative environments for life science research. We thank Shanxi Medical University for the support during the transient translocation period of Zhang’s research group. We would like to thank Yongmei Jiang, Cheng-Yuan Zhang, Dong-Xiao Li, Peng Sun, Yun-Xia Jiang, Shi-Jun Jiang, and Cheng-Mei Li for financial support. Thank Dr. Alexander Zi-Jian Zhang and Emily Zi-Ru Zhang for the final editing and figures. The work is partially supported by NIH grant R01-GM135421 (G.Z.) and by the setup fund from Shanxi Medical University for G.Z.

## Contributions

G.Z. conceived the concepts. L.L. conducted the main experiments. S.M. and G.Z. conducted data analysis. H.L. and Q.Z. contributed to additional data collection. E.G. provided the H3R2me2s antibody and suggestions. X.L. and C.L. generated the JMJD5-depleted A537 cancer cell line. H.L. provided suggestions and final data analysis. G.Z. wrote up the manuscript.

## Supplementary Figures

**sFigure 1.**
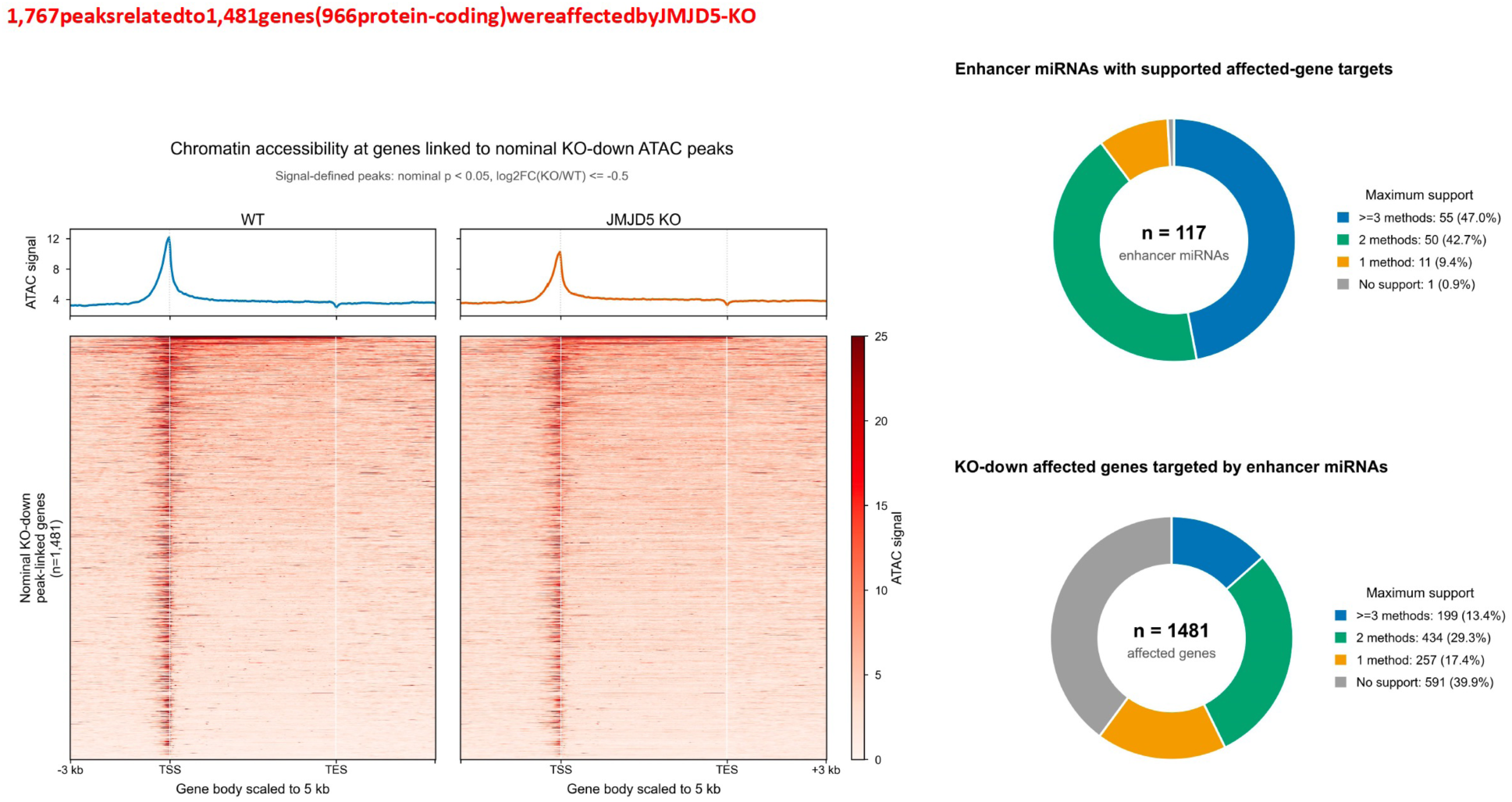
ATAC-seq data show accessibility changes at the transcription start sites of 1,481 genes. Re-analysis of our published ATAC-seq data of mouse embryonic fibroblast cells with and without knockout of JMJD5 shows a decrease in accessibility at the transcription start sites of 1,481 genes with the knockout of JMJD5 (Liu et al., PNAS, 2020). This data is consistent with our current data of CUT&TAG-seq of H3R2me2s in human cancer cells.

**sFigure 2A.**
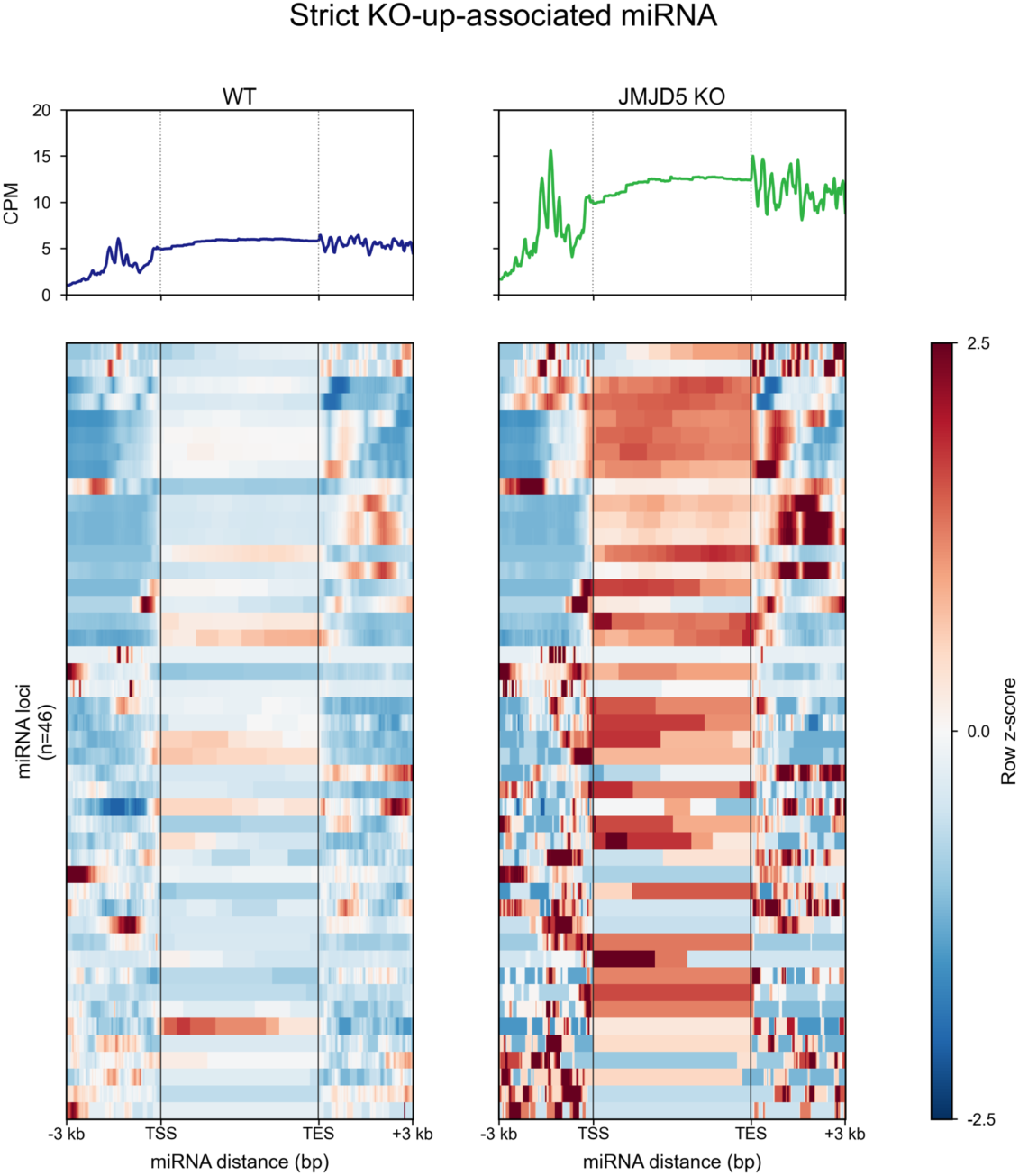
JMJD5 Loss Markedly Increases H3R2me2s across miRNA Loci.

**sFigure 2B.**
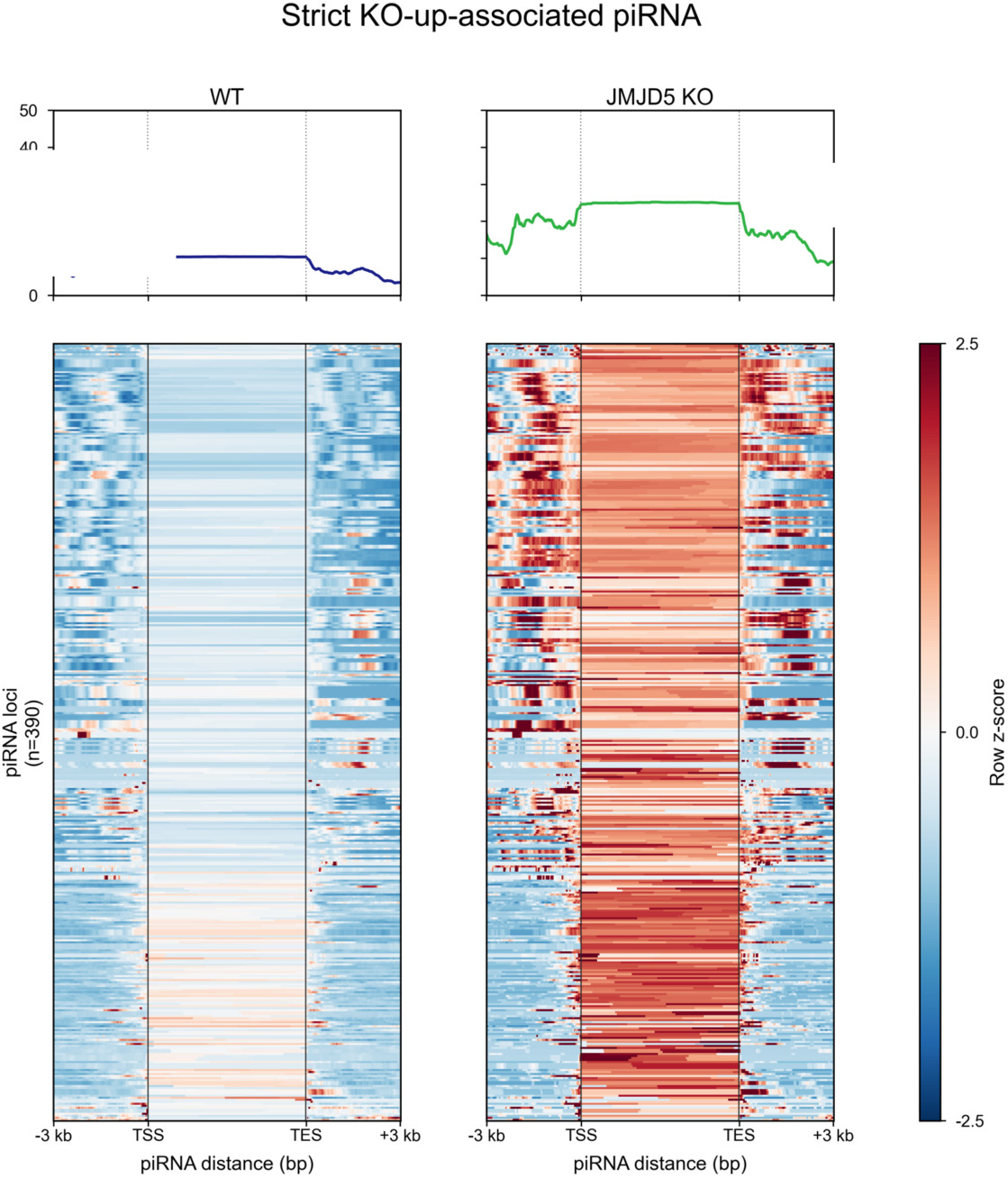
JMJD5 Loss Produces a Pronounced Increase in H3R2me2s across piRNA Loci.

**sFigure 2C.**
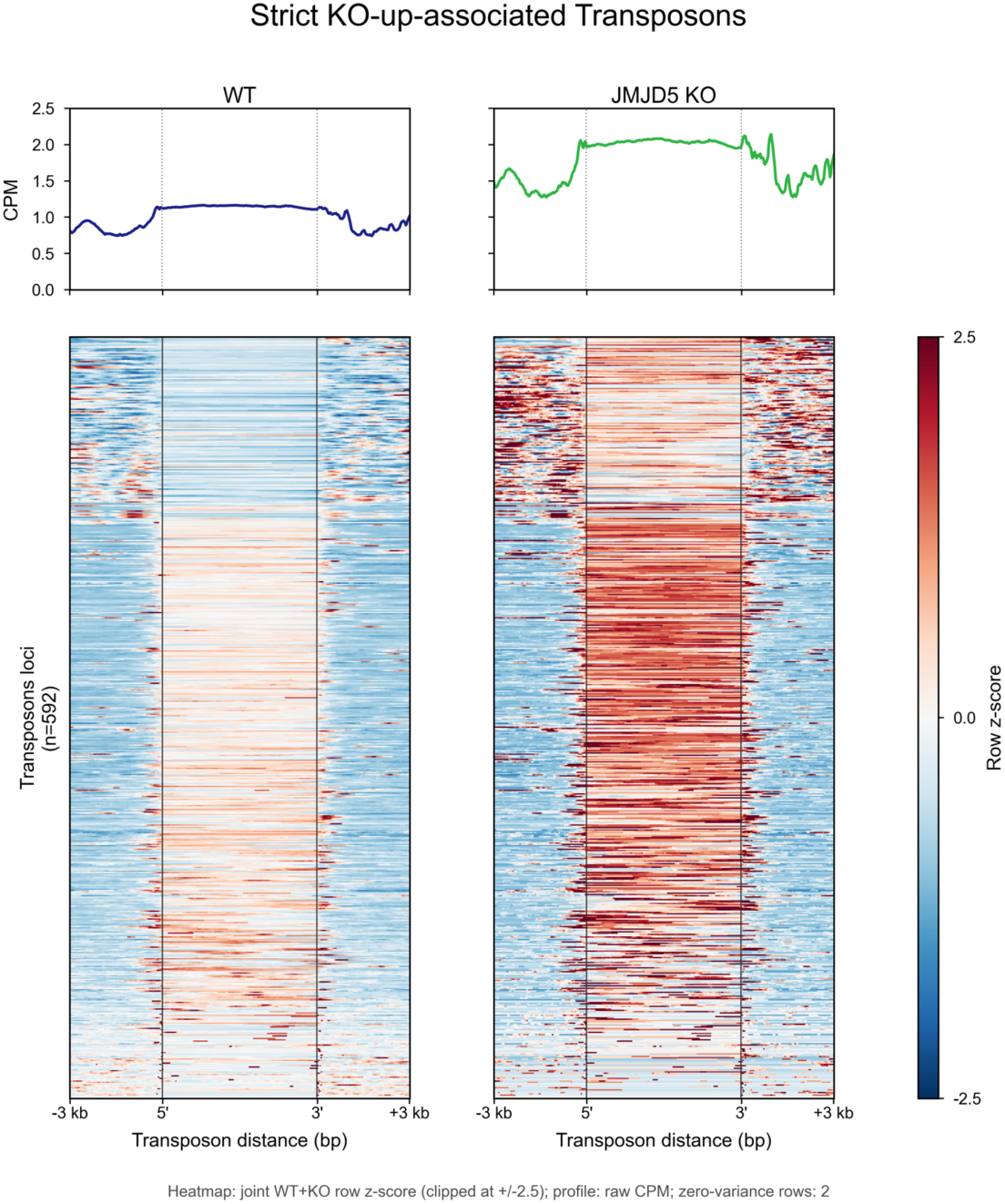
JMJD5 Loss Modestly Elevates H3R2me2s at Transposons, with Enrichment in KO-Up Elements.

**sFigure 2D.**
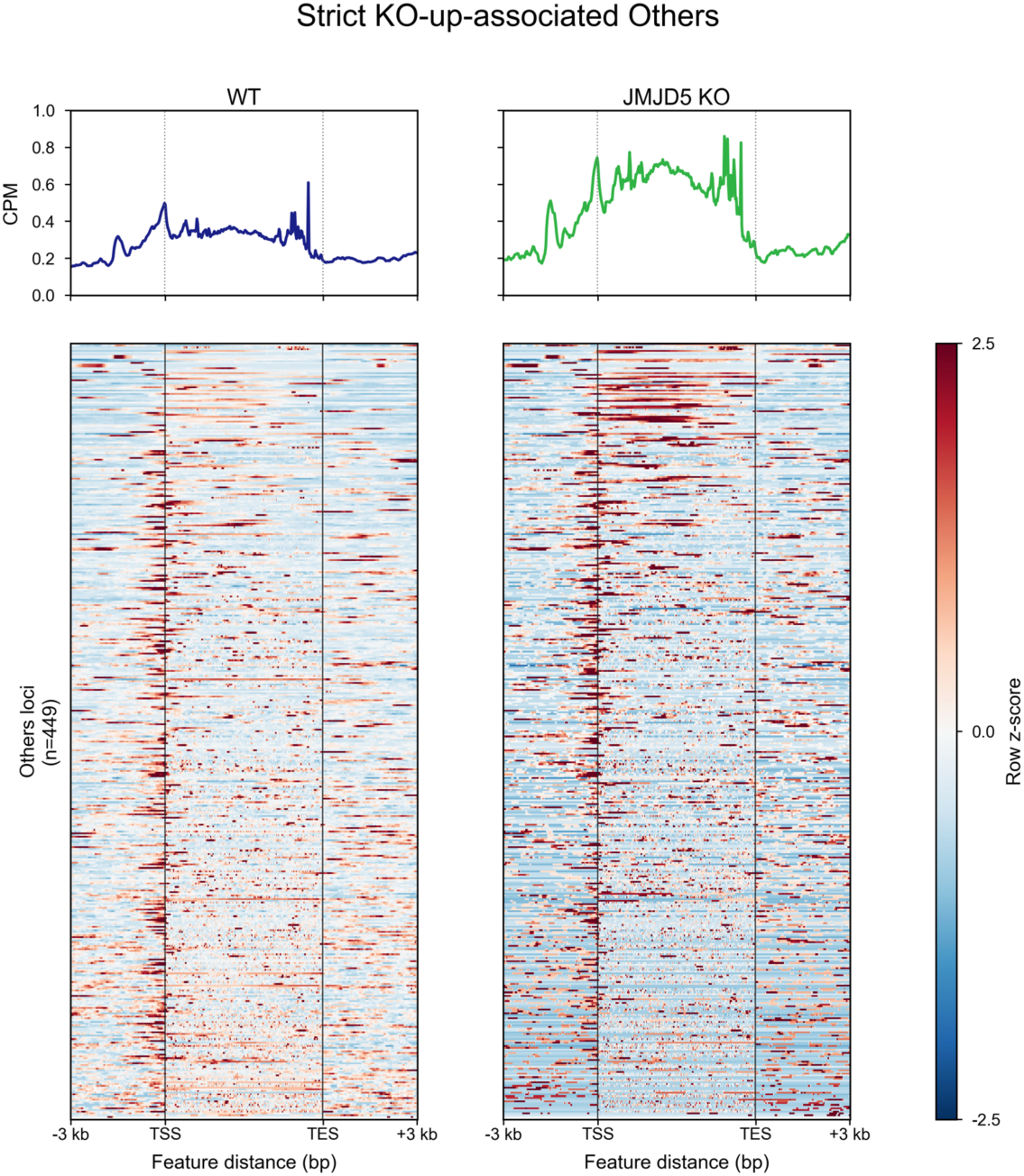
JMJD5 Loss Increases H3R2me2s at Other Non-Coding Genes.

**sFigure 2E.**
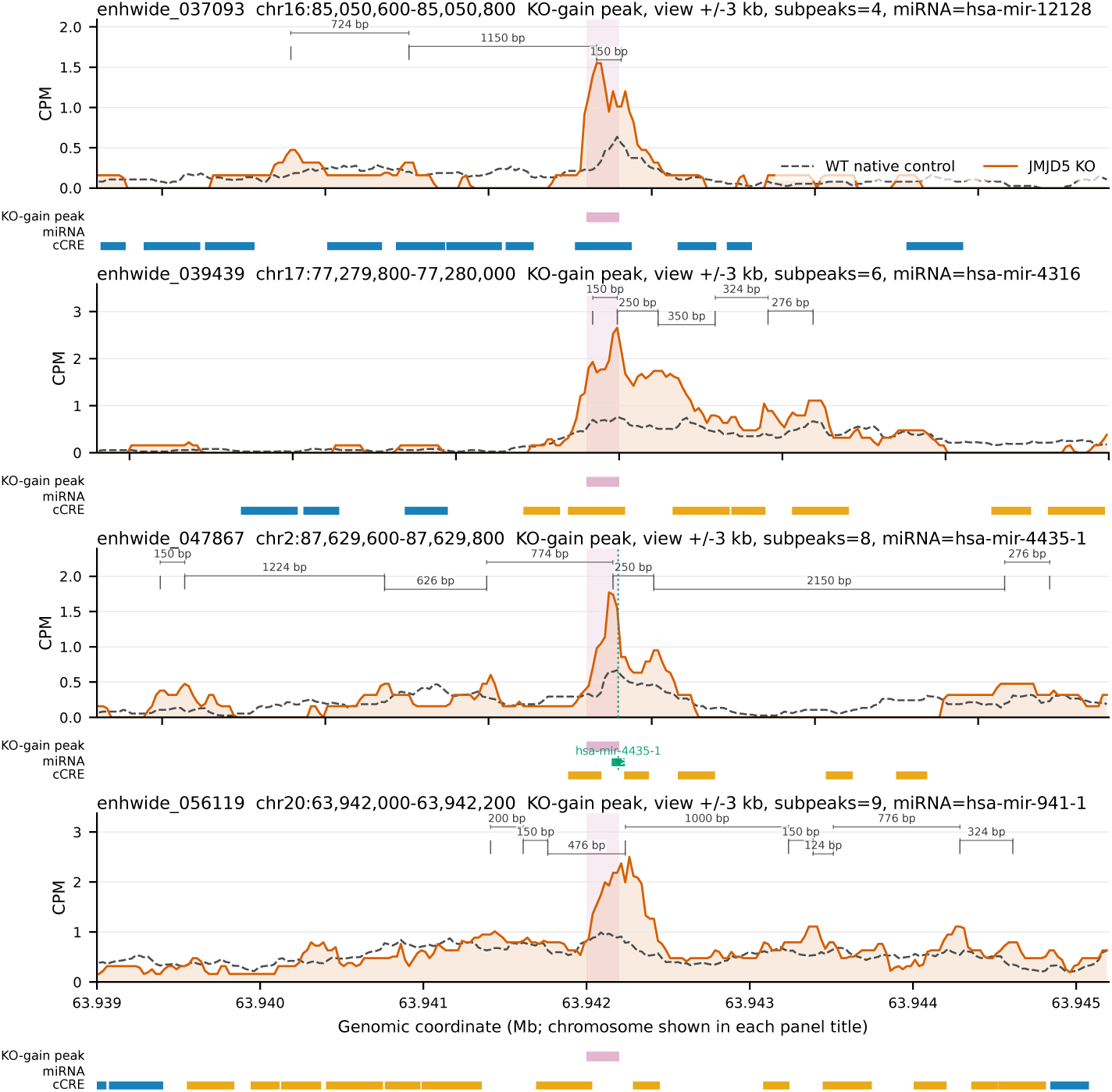
Some rncRNA genes contain two arginine-methylated nucleosomes (+1 and +2) within their gene bodies.

**sFigure 4A, 4B, 4C, 4D, 4E.**
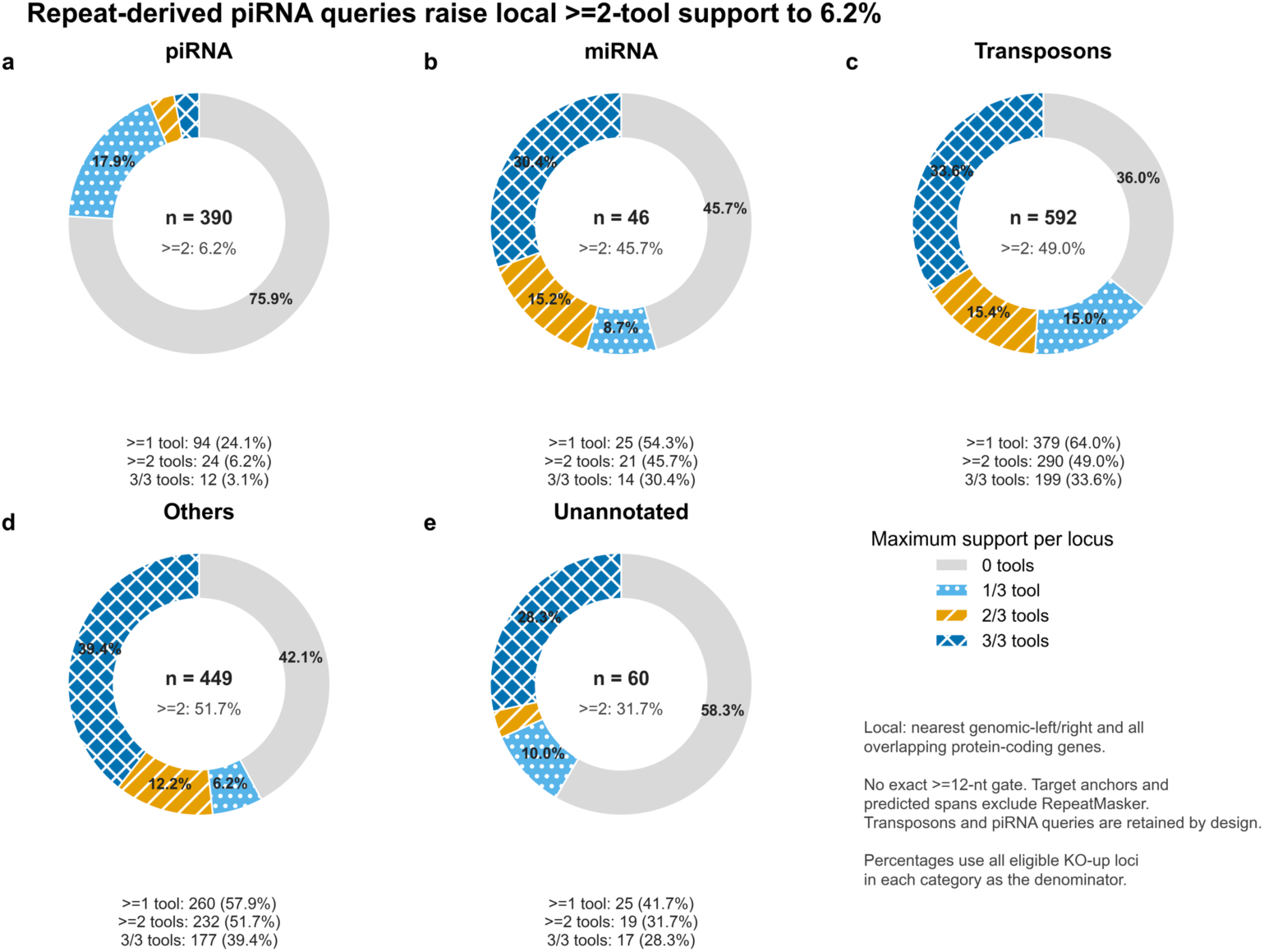
piRNAs, miRNAs, rRNAs, Transponsons, and others were antisense sequences of mRNAs’ 3 ʹ UTRs (regulatory non-coding RNAs, rncRNAs).

**sFigure 4E, 4G.**
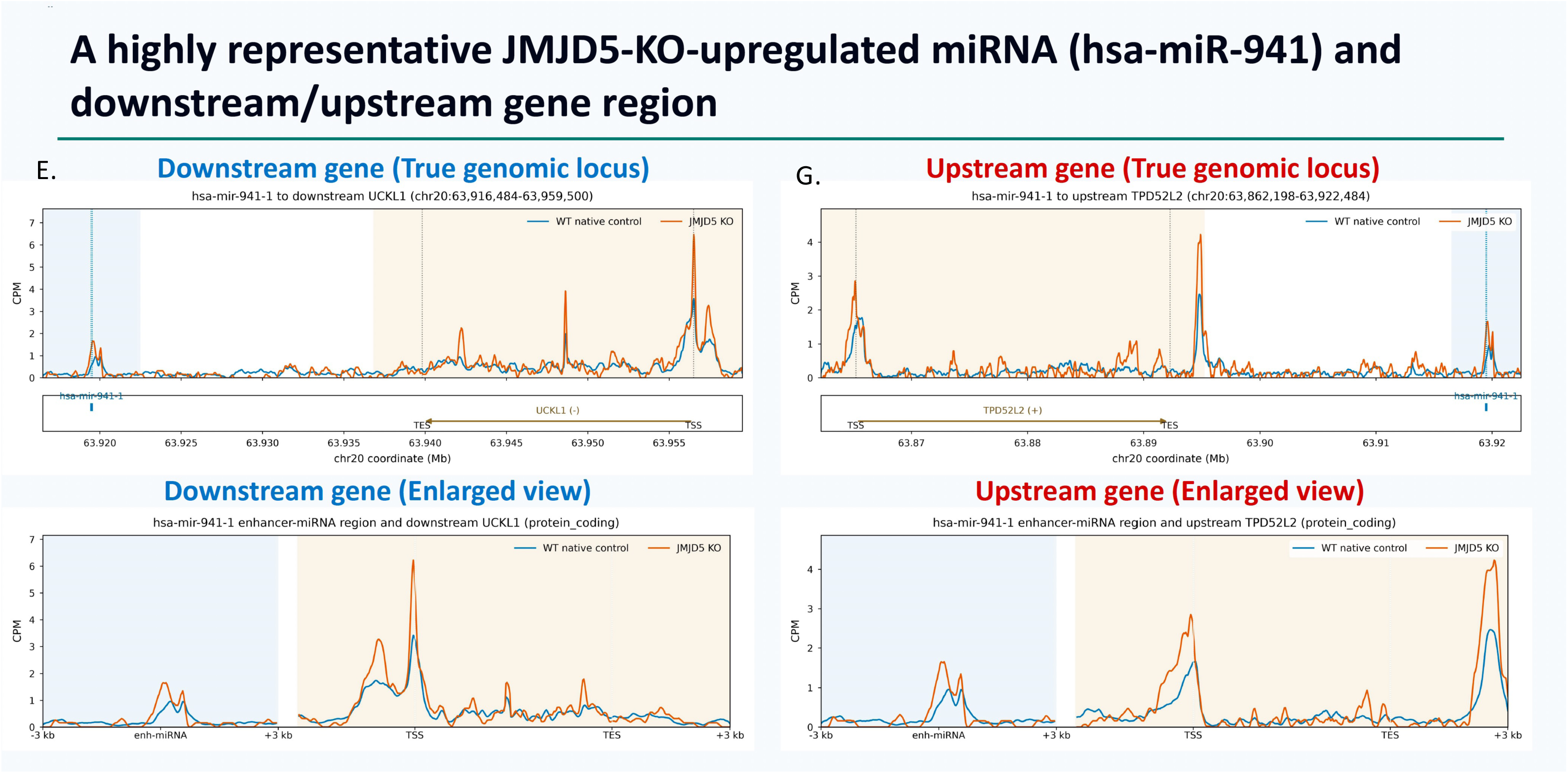
Has-miR-941 is located downstream of its two target genes.

**sFigure 4H.**
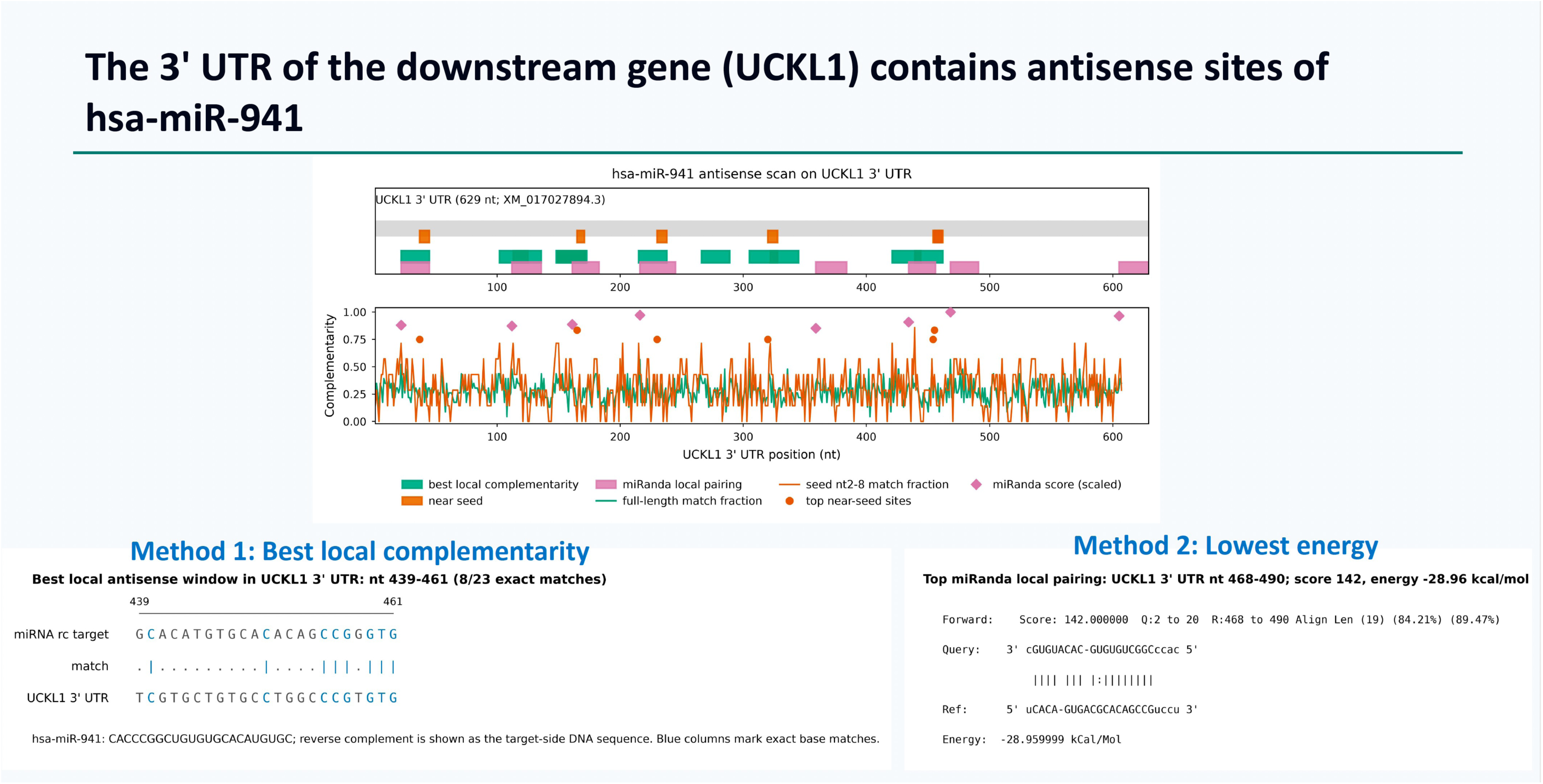
The match between has-miR-941 and the 3 ’ UTR of the UCKL1 gene.

**sFigure 4I.**
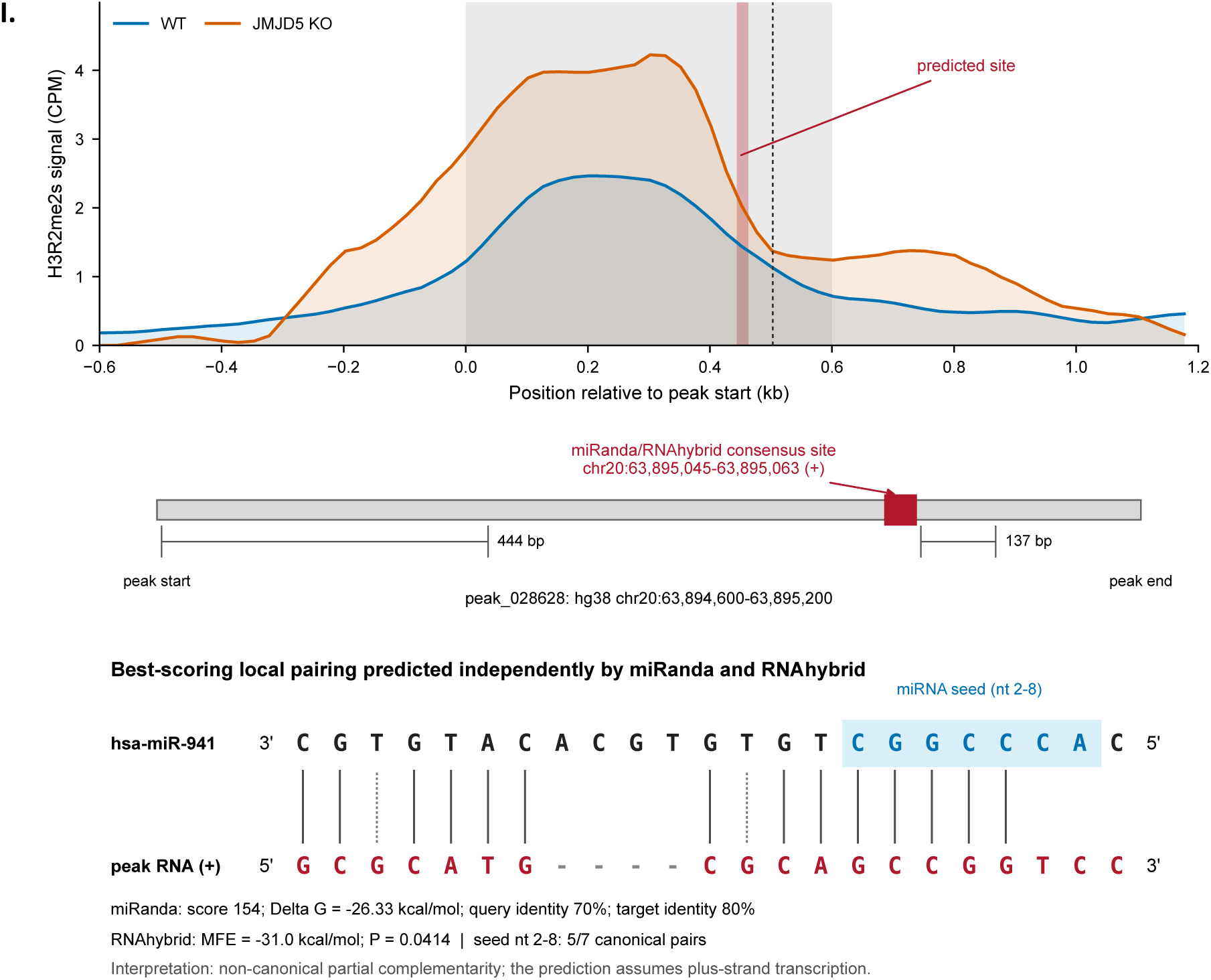
The match between has-miR-941 and the 3 ’ UTR of the TPD52L1 gene.

**sFigure 5A, 5B.**
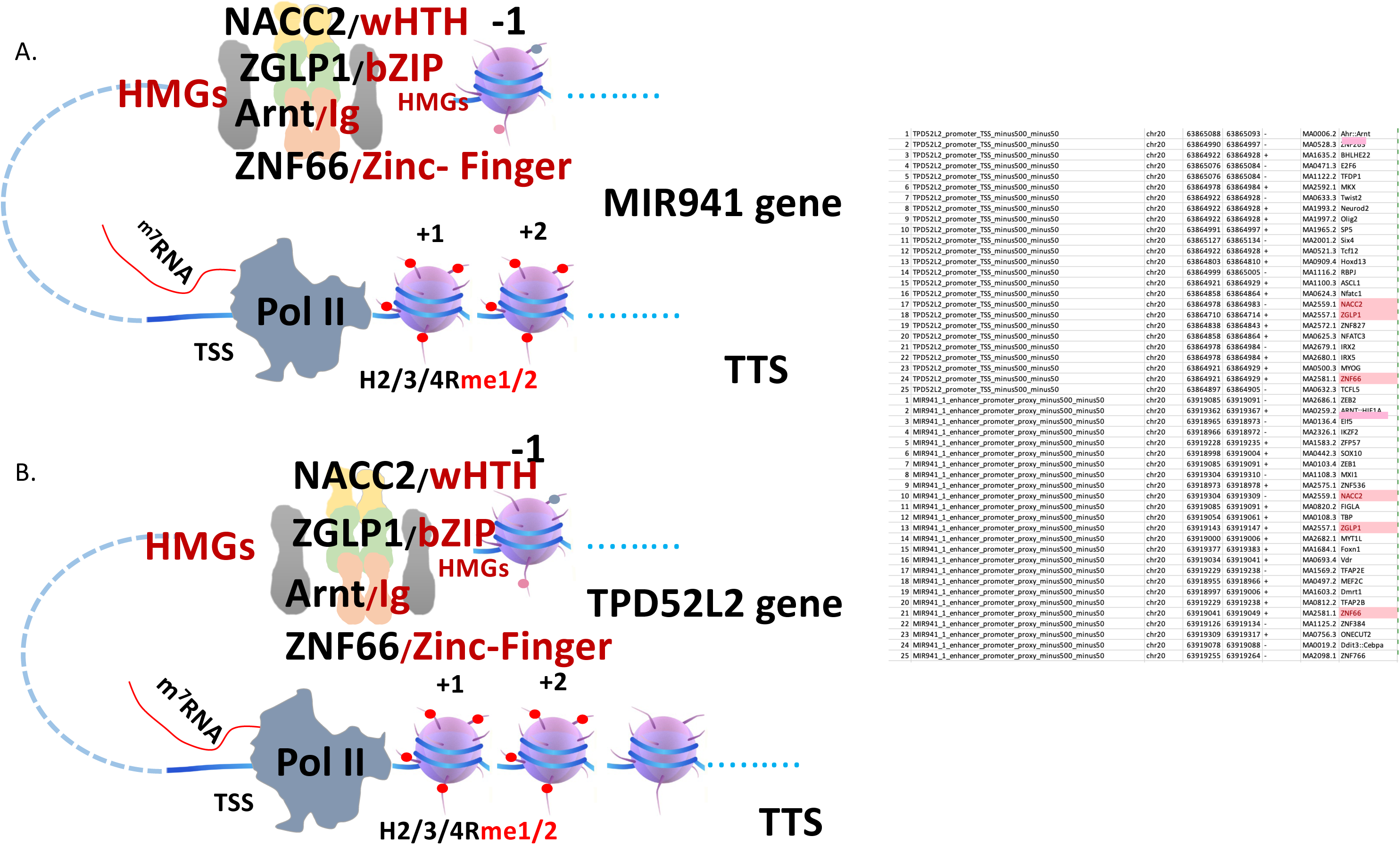
rncRNA and mRNA share transcription factors.

## Reference

An, W., J. Kim and R. G. Roeder (2004). “Ordered cooperative functions of PRMT1, p300, and CARM1 in transcriptional activation by p53.” Cell 117(6): 735–748.

Bachand, F. (2007). “Protein arginine methyltransferases: from unicellular eukaryotes to humans.” Eukaryot Cell 6(6): 889–898.

Banerji, J., L. Olson and W. Schaffner (1983). “A lymphocyte-specific cellular enhancer is located downstream of the joining region in immunoglobulin heavy chain genes.” Cell 33(3): 729–740.

Banerji, J., S. Rusconi and W. Schaffner (1981). “Expression of a beta-globin gene is enhanced by remote SV40 DNA sequences.” Cell 27(2 Pt 1): 299–308.

Barckmann, B., S. Pierson, J. Dufourt, C. Papin, C. Armenise, F. Port, T. Grentzinger, S. Chambeyron, G. Baronian, J. P. Desvignes, T. Curk and M. Simonelig (2015). “Aubergine iCLIP Reveals piRNA-Dependent Decay of mRNAs Involved in Germ Cell Development in the Early Embryo.” Cell Rep 12(7): 1205–1216.

Batut, P. J., X. Y. Bing, Z. Sisco, J. Raimundo, M. Levo and M. S. Levine (2022). “Genome organization controls transcriptional dynamics during development.” Science 375(6580): 566–570.

Benoist, C. and P. Chambon (1981). “In vivo sequence requirements of the SV40 early promotor region.” Nature 290(5804): 304–310.

Cai, X., C. H. Hagedorn and B. R. Cullen (2004). “Human microRNAs are processed from capped, polyadenylated transcripts that can also function as mRNAs.” RNA 10(12): 1957–1966.

Carmell, M. A., A. Girard, H. J. van de Kant, D. Bourc’his, T. H. Bestor, D. G. de Rooij and G. J. Hannon (2007). “MIWI2 is essential for spermatogenesis and repression of transposons in the mouse male germline.” Dev Cell 12(4): 503–514.

Cecchini, K., M. Zamani, N. Ajaykumar, J. Vega-Badillo, A. Bagci, S. Bailey, P. D. Zamore and I. Gainetdinov (2026). “Cleavage of mRNAs by a minority of pachytene piRNAs improves sperm fitness.” Nature 652(8109): 508–516.

Chen, D., S. M. Huang and M. R. Stallcup (2000). “Synergistic, p160 coactivator-dependent enhancement of estrogen receptor function by CARM1 and p300.” J Biol Chem 275(52): 40810–40816.

Chung, W. J., K. Okamura, R. Martin and E. C. Lai (2008). “Endogenous RNA interference provides a somatic defense against Drosophila transposons.” Curr Biol 18(11): 795–802.

Cox, D. N., A. Chao, J. Baker, L. Chang, D. Qiao and H. Lin (1998). “A novel class of evolutionarily conserved genes defined by piwi are essential for stem cell self-renewal.” Genes Dev 12(23): 3715–3727.

Daujat, S., U. M. Bauer, V. Shah, B. Turner, S. Berger and T. Kouzarides (2002). “Crosstalk between CARM1 methylation and CBP acetylation on histone H3.” Curr Biol 12(24): 2090–2097.

De Santa, F., I. Barozzi, F. Mietton, S. Ghisletti, S. Polletti, B. K. Tusi, H. Muller, J. Ragoussis, C. L. Wei and G. Natoli (2010). “A large fraction of extragenic RNA pol II transcription sites overlap enhancers.” PLoS Biol 8(5): e1000384.

Deng, W. and H. Lin (2002). “miwi, a murine homolog of piwi, encodes a cytoplasmic protein essential for spermatogenesis.” Dev Cell 2(6): 819–830.

Dixon, J. R., S. Selvaraj, F. Yue, A. Kim, Y. Li, Y. Shen, M. Hu, J. S. Liu and B. Ren (2012). “Topological domains in mammalian genomes identified by analysis of chromatin interactions.” Nature 485(7398): 376–380.

Eulalio, A., E. Huntzinger, T. Nishihara, J. Rehwinkel, M. Fauser and E. Izaurralde (2009). “Deadenylation is a widespread effect of miRNA regulation.” RNA 15(1): 21–32.

Fire, A., S. Xu, M. K. Montgomery, S. A. Kostas, S. E. Driver and C. C. Mello (1998). “Potent and specific genetic interference by double-stranded RNA in Caenorhabditis elegans.” Nature 391(6669): 806–811.

Foster, C. T., O. M. Dovey, L. Lezina, J. L. Luo, T. W. Gant, N. Barlev, A. Bradley and S. M. Cowley (2010). “Lysine-specific demethylase 1 regulates the embryonic transcriptome and CoREST stability.” Mol Cell Biol 30(20): 4851–4863.

Giese, K., C. Kingsley, J. R. Kirshner and R. Grosschedl (1995). “Assembly and function of a TCR alpha enhancer complex is dependent on LEF-1-induced DNA bending and multiple protein-protein interactions.” Genes Dev 9(8): 995–1008.

Gillies, S. D., S. L. Morrison, V. T. Oi and S. Tonegawa (1983). “A tissue-specific transcription enhancer element is located in the major intron of a rearranged immunoglobulin heavy chain gene.” Cell 33(3): 717–728.

Goh, W. S., I. Falciatori, O. H. Tam, R. Burgess, O. Meikar, N. Kotaja, M. Hammell and G. J. Hannon (2015). “piRNA-directed cleavage of meiotic transcripts regulates spermatogenesis.” Genes Dev 29(10): 1032–1044.

Gou, L. T., P. Dai, J. H. Yang, Y. Xue, Y. P. Hu, Y. Zhou, J. Y. Kang, X. Wang, H. Li, M. M. Hua, S. Zhao, S. D. Hu, L. G. Wu, H. J. Shi, Y. Li, X. D. Fu, L. H. Qu, E. D. Wang and M. F. Liu (2014). “Pachytene piRNAs instruct massive mRNA elimination during late spermiogenesis.” Cell Res 24(6): 680–700.

Grosschedl, R. and M. L. Birnstiel (1980). “Spacer DNA sequences upstream of the T-A-T-A-A-A-T-A sequence are essential for promotion of H2A histone gene transcription in vivo.” Proc Natl Acad Sci U S A 77(12): 7102–7106.

Gruss, P., R. Dhar and G. Khoury (1981). “Simian virus 40 tandem repeated sequences as an element of the early promoter.” Proc Natl Acad Sci U S A 78(2): 943–947.

Guccione, E., C. Bassi, F. Casadio, F. Martinato, M. Cesaroni, H. Schuchlautz, B. Luscher and B. Amati (2007). “Methylation of histone H3R2 by PRMT6 and H3K4 by an MLL complex are mutually exclusive.” Nature 449(7164): 933–937.

Gunawardane, L. S., K. Saito, K. M. Nishida, K. Miyoshi, Y. Kawamura, T. Nagami, H. Siomi and M. C. Siomi (2007). “A slicer-mediated mechanism for repeat-associated siRNA 5’ end formation in Drosophila.” Science 315(5818): 1587–1590.

Hale, B. G. (2022). “Antiviral immunity triggered by infection-induced host transposable elements.” Curr Opin Virol 52: 211–216.

Hamilton, A. J. and D. C. Baulcombe (1999). “A species of small antisense RNA in posttranscriptional gene silencing in plants.” Science 286(5441): 950–952.

Han, J., Y. Lee, K. H. Yeom, Y. K. Kim, H. Jin and V. N. Kim (2004). “The Drosha-DGCR8 complex in primary microRNA processing.” Genes Dev 18(24): 3016–3027.

Hawkins, P. G. and K. V. Morris (2008). “RNA and transcriptional modulation of gene expression.” Cell Cycle 7(5): 602–607.

Hnisz, D., B. J. Abraham, T. I. Lee, A. Lau, V. Saint-Andre, A. A. Sigova, H. A. Hoke and R. A. Young (2013). “Super-enhancers in the control of cell identity and disease.” Cell 155(4): 934–947.

Hong, X., J. Zang, J. White, C. Wang, C. H. Pan, R. Zhao, R. C. Murphy, S. Dai, P. Henson, J. W. Kappler, J. Hagman and G. Zhang (2010). “Interaction of JMJD6 with single-stranded RNA.” Proc Natl Acad Sci U S A 107(33): 14568–14572.

Houwing, S., L. M. Kamminga, E. Berezikov, D. Cronembold, A. Girard, H. van den Elst, D. V. Filippov, H. Blaser, E. Raz, C. B. Moens, R. H. Plasterk, G. J. Hannon, B. W. Draper and R. F. Ketting (2007). “A role for Piwi and piRNAs in germ cell maintenance and transposon silencing in Zebrafish.” Cell 129(1): 69–82.

Kawasaki, H. and K. Taira (2004). “MicroRNA-196 inhibits HOXB8 expression in myeloid differentiation of HL60 cells.” Nucleic Acids Symp Ser (Oxf)(48): 211–212.

Kaya-Okur, H. S., S. J. Wu, C. A. Codomo, E. S. Pledger, T. D. Bryson, J. G. Henikoff, K. Ahmad and S. Henikoff (2019). “CUT&Tag for efficient epigenomic profiling of small samples and single cells.” Nat Commun 10(1): 1930.

Kim, T. K., M. Hemberg, J. M. Gray, A. M. Costa, D. M. Bear, J. Wu, D. A. Harmin, M. Laptewicz, K. Barbara-Haley, S. Kuersten, E. Markenscoff-Papadimitriou, D. Kuhl, H. Bito, P. F. Worley, G. Kreiman and M. E. Greenberg (2010). “Widespread transcription at neuronal activity-regulated enhancers.” Nature 465(7295): 182–187.

Larsen, S. C., K. B. Sylvestersen, A. Mund, D. Lyon, M. Mullari, M. V. Madsen, J. A. Daniel, L. J. Jensen and M. L. Nielsen (2016). “Proteome-wide analysis of arginine monomethylation reveals widespread occurrence in human cells.” Sci Signal 9(443): rs9.

Lee, R. C., R. L. Feinbaum and V. Ambros (1993). “The C. elegans heterochronic gene lin-4 encodes small RNAs with antisense complementarity to lin-14.” Cell 75(5): 843–854.

Lee, S., H. Liu, R. Hill, C. Chen, X. Hong, F. Crawford, M. Kingsley, Q. Zhang, X. Liu, Z. Chen, A. Lengeling, K. M. Bernt, P. Marrack, J. Kappler, Q. Zhou, C. Y. Li, Y. Xue, K. Hansen and G. Zhang (2020). “JMJD6 cleaves MePCE to release positive transcription elongation factor b (P-TEFb) in higher eukaryotes.” Elife 9.

Lee, Y., C. Ahn, J. Han, H. Choi, J. Kim, J. Yim, J. Lee, P. Provost, O. Radmark, S. Kim and V. N. Kim (2003). “The nuclear RNase III Drosha initiates microRNA processing.” Nature 425(6956): 415–419.

Lee, Y., M. Kim, J. Han, K. H. Yeom, S. Lee, S. H. Baek and V. N. Kim (2004). “MicroRNA genes are transcribed by RNA polymerase II.” EMBO J 23(20): 4051–4060.

Lee, Y., A. J. Song, R. Baker, B. Micales, S. J. Conway and G. E. Lyons (2000). “Jumonji, a nuclear protein that is necessary for normal heart development.” Circ Res 86(9): 932–938.

Li, L., S. Hunter, S. Leach, Y. Zhuang, H. Liu, J. Gao, Q. Zhang, T. J. Stasevich, H. Kimura, R. Dowell and G. Zhang (2025). “Fragile nucleosomes are essential for RNA Polymerase II to transcribe in eukaryotes.” bioRxiv.

Li, X., X. Tang, X. Bing, C. Catalano, T. Li, G. Dolsten, C. Wu and M. Levine (2023). “GAGA-associated factor fosters loop formation in the Drosophila genome.” Mol Cell 83(9): 1519–1526 e1514.

Liang, Y., Q. Zou and W. Yu (2017). “Steering Against Wind: A New Network of NamiRNAs and Enhancers.” Genomics Proteomics Bioinformatics 15(5): 331–337.

Liu, H., S. Lee, Q. Zhang, Z. Chen and G. Zhang (2020). “The potential underlying mechanism of the leukemia caused by MLL-fusion and potential treatments.” Mol Carcinog.

Liu, H., L. Li, J. Gao, S. Leach, P. Wei, Q. Zhang, H. Huang, T. Houles, P. P. Roux and G. Zhang (2025). “CDK11 activates CDK12 to trigger the elongation of RNA Polymerase II.” bioRxiv.

Liu, H., L. Li, Q. Zhang, C. Ott and G. Zhang (2025). “Enhancer/Enhanceosome/Super-enhancer and the promoter-proximate RNA Polymerase II (Pol II) pausing are coupled.” bioRxiv.

Liu, H., Li, L., Zhang, Q., Ott, C., Zhang, G. (2025). “Enhancer/Enhanceosome/Super-enhancer and the promoter-proximal RNA Polymerase II (Pol II) pausing are coupled.” BioRxiv.

Liu, H., S. Ramachandran, N. Fong, T. Phang, S. Lee, P. Parsa, X. Liu, L. Harmacek, T. Danhorn, T. Song, S. Oh, Q. Zhang, Z. Chen, Q. Zhang, T. H. Tu, C. Happoldt, B. O’Conner, R. Janknecht, C. Y. Li, P. Marrack, J. Kappler, S. Leach and G. Zhang (2020). “JMJD5 couples with CDK9 to release the paused RNA polymerase II.” Proc Natl Acad Sci U S A 117(33): 19888–19895.

Liu, H., C. Wang, S. Lee, Y. Deng, M. Wither, S. Oh, F. Ning, C. Dege, Q. Zhang, X. Liu, A. M. Johnson, J. Zang, Z. Chen, R. Janknecht, K. Hansen, P. Marrack, C. Y. Li, J. W. Kappler, J. Hagman and G. Zhang (2017). “Clipping of arginine-methylated histone tails by JMJD5 and JMJD7.” Proc Natl Acad Sci U S A 114(37): E7717–E7726.

Liu, H., C. Wang, S. Lee, F. Ning, Y. Wang, Q. Zhang, Z. Chen, J. Zang, J. Nix, S. Dai, P. Marrack, J. Hagman, J. Kappler and G. Zhang (2018). “Specific Recognition of Arginine Methylated Histone Tails by JMJD5 and JMJD7.” Sci Rep 8(1): 3275.

Liu, H., P. Wei, Q. Zhang, Z. Chen, J. Liu and G. Zhang (2022). “The Novel Protease Activities of JMJD5-JMJD6-JMJD7 and Arginine Methylation Activities of Arginine Methyltransferases Are Likely Coupled.” Biomolecules 12(3).

Liu, Y. P., J. Haasnoot, O. ter Brake, B. Berkhout and P. Konstantinova (2008). “Inhibition of HIV-1 by multiple siRNAs expressed from a single microRNA polycistron.” Nucleic Acids Res 36(9): 2811–2824.

Ma, H., C. T. Baumann, H. Li, B. D. Strahl, R. Rice, M. A. Jelinek, D. W. Aswad, C. D. Allis, G. L. Hager and M. R. Stallcup (2001). “Hormone-dependent, CARM1-directed, arginine-specific methylation of histone H3 on a steroid-regulated promoter.” Curr Biol 11(24): 1981–1985.

Meister, G., M. Landthaler, A. Patkaniowska, Y. Dorsett, G. Teng and T. Tuschl (2004). “Human Argonaute2 mediates RNA cleavage targeted by miRNAs and siRNAs.” Mol Cell 15(2): 185–197.

Mercola, M., X. F. Wang, J. Olsen and K. Calame (1983). “Transcriptional enhancer elements in the mouse immunoglobulin heavy chain locus.” Science 221(4611): 663–665.

Migliori, V., J. Muller, S. Phalke, D. Low, M. Bezzi, W. C. Mok, S. K. Sahu, J. Gunaratne, P. Capasso, C. Bassi, V. Cecatiello, A. De Marco, W. Blackstock, V. Kuznetsov, B. Amati, M. Mapelli and E. Guccione (2012). “Symmetric dimethylation of H3R2 is a newly identified histone mark that supports euchromatin maintenance.” Nat Struct Mol Biol 19(2): 136–144.

Moore, L. D., T. Le and G. Fan (2013). “DNA methylation and its basic function.” Neuropsychopharmacology 38(1): 23–38.

Nora, E. P., B. R. Lajoie, E. G. Schulz, L. Giorgetti, I. Okamoto, N. Servant, T. Piolot, N. L. van Berkum, J. Meisig, J. Sedat, J. Gribnau, E. Barillot, N. Bluthgen, J. Dekker and E. Heard (2012). “Spatial partitioning of the regulatory landscape of the X-inactivation centre.” Nature 485(7398): 381–385.

Oh, S. and R. Janknecht (2012). “Histone demethylase JMJD5 is essential for embryonic development.” Biochem Biophys Res Commun 420(1): 61–65.

Pal, S., S. N. Vishwanath, H. Erdjument-Bromage, P. Tempst and S. Sif (2004). “Human SWI/SNF-associated PRMT5 methylates histone H3 arginine 8 and negatively regulates expression of ST7 and NM23 tumor suppressor genes.” Mol Cell Biol 24(21): 9630–9645.

Parker, S. C., M. L. Stitzel, D. L. Taylor, J. M. Orozco, M. R. Erdos, J. A. Akiyama, K. L. van Bueren, P. S. Chines, N. Narisu, N. C. S. Program, B. L. Black, A. Visel, L. A. Pennacchio, F. S. Collins, A. National Institutes of Health Intramural Sequencing Center Comparative Sequencing Program and N. C. S. P. Authors (2013). “Chromatin stretch enhancer states drive cell-specific gene regulation and harbor human disease risk variants.” Proc Natl Acad Sci U S A 110(44): 17921–17926.

Pollack, B. P., S. V. Kotenko, W. He, L. S. Izotova, B. L. Barnoski and S. Pestka (1999). “The human homologue of the yeast proteins Skb1 and Hsl7p interacts with Jak kinases and contains protein methyltransferase activity.” J Biol Chem 274(44): 31531–31542.

Rakow, S., S. S. Pullamsetti, U. M. Bauer and C. Bouchard (2020). “Assaying epigenome functions of PRMTs and their substrates.” Methods 175: 53–65.

Ramirez, F., D. P. Ryan, B. Gruning, V. Bhardwaj, F. Kilpert, A. S. Richter, S. Heyne, F. Dundar and T. Manke (2016). “deepTools2: a next generation web server for deep-sequencing data analysis.” Nucleic Acids Res 44(W1): W160–165.

Ren, X., G. Liu and J. Zhou (2025). “Nuclear-activating miRNAs: unveiling the intricacies of subcellular miRNA function and regulation in cancer and immunity disease.” Cancer Cell Int 25(1): 147.

Rodriguez, A., S. Griffiths-Jones, J. L. Ashurst and A. Bradley (2004). “Identification of mammalian microRNA host genes and transcription units.” Genome Res 14(10A): 1902–1910.

Ross-Innes, C. S., R. Stark, A. E. Teschendorff, K. A. Holmes, H. R. Ali, M. J. Dunning, G. D. Brown, O. Gojis, I. O. Ellis, A. R. Green, S. Ali, S. F. Chin, C. Palmieri, C. Caldas and J. S. Carroll (2012). “Differential oestrogen receptor binding is associated with clinical outcome in breast cancer.” Nature 481(7381): 389–393.

Scaglione, A., J. Patzig, J. Liang, R. Frawley, J. Bok, A. Mela, C. Yattah, J. Zhang, S. X. Teo, T. Zhou, S. Chen, E. Bernstein, P. Canoll, E. Guccione and P. Casaccia (2018). “PRMT5-mediated regulation of developmental myelination.” Nat Commun 9(1): 2840.

Strahl, B. D., S. D. Briggs, C. J. Brame, J. A. Caldwell, S. S. Koh, H. Ma, R. G. Cook, J. Shabanowitz, D. F. Hunt, M. R. Stallcup and C. D. Allis (2001). “Methylation of histone H4 at arginine 3 occurs in vivo and is mediated by the nuclear receptor coactivator PRMT1.” Curr Biol 11(12): 996–1000.

Tan, Y., B. Zhang, T. Wu, G. Skogerbo, X. Zhu, X. Guo, S. He and R. Chen (2009). “Transcriptional inhibiton of Hoxd4 expression by miRNA-10a in human breast cancer cells.” BMC Mol Biol 10: 12.

Tang, S. C., U. Vijayakumar, Y. Zhang and M. J. Fullwood (2022). “Super-Enhancers, Phase-Separated Condensates, and 3D Genome Organization in Cancer.” Cancers (Basel) 14(12).

Thanos, D. and T. Maniatis (1995). “Virus induction of human IFN beta gene expression requires the assembly of an enhanceosome.” Cell 83(7): 1091–1100.

Waldmann, T., A. Izzo, K. Kamieniarz, F. Richter, C. Vogler, B. Sarg, H. Lindner, N. L. Young, G. Mittler, B. A. Garcia and R. Schneider (2011). “Methylation of H2AR29 is a novel repressive PRMT6 target.” Epigenetics Chromatin 4: 11.

Wang, X. J., J. L. Reyes, N. H. Chua and T. Gaasterland (2004). “Prediction and identification of Arabidopsis thaliana microRNAs and their mRNA targets.” Genome Biol 5(9): R65.

Watanabe, T., E. C. Cheng, M. Zhong and H. Lin (2015). “Retrotransposons and pseudogenes regulate mRNAs and lncRNAs via the piRNA pathway in the germline.” Genome Res 25(3): 368–380.

Whyte, W. A., D. A. Orlando, D. Hnisz, B. J. Abraham, C. Y. Lin, M. H. Kagey, P. B. Rahl, T. I. Lee and R. A. Young (2013). “Master transcription factors and mediator establish super-enhancers at key cell identity genes.” Cell 153(2): 307–319.

Wightman, B., I. Ha and G. Ruvkun (1993). “Posttranscriptional regulation of the heterochronic gene lin-14 by lin-4 mediates temporal pattern formation in C. elegans.” Cell 75(5): 855–862.

Williams, A. E. (2008). “Functional aspects of animal microRNAs.” Cell Mol Life Sci 65(4): 545–562.

Xiao, M., J. Li, W. Li, Y. Wang, F. Wu, Y. Xi, L. Zhang, C. Ding, H. Luo, Y. Li, L. Peng, L. Zhao, S. Peng, Y. Xiao, S. Dong, J. Cao and W. Yu (2017). “MicroRNAs activate gene transcription epigenetically as an enhancer trigger.” RNA Biol 14(10): 1326–1334.

Yin, H. and H. Lin (2007). “An epigenetic activation role of Piwi and a Piwi-associated piRNA in Drosophila melanogaster.” Nature 450(7167): 304–308.

Zamore, P. D., T. Tuschl, P. A. Sharp and D. P. Bartel (2000). “RNAi: double-stranded RNA directs the ATP-dependent cleavage of mRNA at 21 to 23 nucleotide intervals.” Cell 101(1): 25–33.

Zhang, G. (2024). “A hypothetical model: Chromatin remodelers couple with acetyltransferases to trigger the elongation of RNA polymerase II (pol II).” Front Epigenet Epigenom 2.

Zhang, G. (2026). “DSIF, NELF, the Integrator complex, and the promoter-proximal RNA Polymerase II pausing are coupled in metazoans.” Gene 1015: 150422.

Zhang, N., Z. Fu, S. Linke, J. Chicher, J. J. Gorman, D. Visk, G. G. Haddad, L. Poellinger, D. J. Peet, F. Powell and R. S. Johnson (2010). “The asparaginyl hydroxylase factor inhibiting HIF-1alpha is an essential regulator of metabolism.” Cell Metab 11(5): 364–378.

Zhang, P., J. Y. Kang, L. T. Gou, J. Wang, Y. Xue, G. Skogerboe, P. Dai, D. W. Huang, R. Chen, X. D. Fu, M. F. Liu and S. He (2015). “MIWI and piRNA-mediated cleavage of messenger RNAs in mouse testes.” Cell Res 25(2): 193–207.

Zheng, G., J. A. Dahl, Y. Niu, Y. Fu, A. Klungland, Y. G. Yang and C. He (2013). “Sprouts of RNA epigenetics: the discovery of mammalian RNA demethylases.” RNA Biol 10(6): 915–918.

